# Genome-wide analysis of the non-coding RNA synthetic genetic network reveals extensive plasticity and distinct environmental dependent roles for U3 snoRNA paralogs

**DOI:** 10.1101/2022.03.10.483001

**Authors:** Soukaina Timouma, Tanda Qi, Marcin G Fraczek, Laura Natalia Balarezo-Cisneros, Steven Parker, Kobchai Duangrattanalert, Matthew Wenjie Feng, Sam Griffiths-Jones, Raymond T O’Keefe, Daniela Delneri

## Abstract

Non-coding RNAs (ncRNAs) with unknown function have been found to be responsible for fitness changes in yeast. With the help of synthetic genetic array (SGA) methodology, genetic interaction screenings between protein-coding genes have contributed to the understanding of complex genetic pathways in eukaryotes. By extending this approach to the ncRNAs, we created more than 15,000 ncRNA double mutants and scored their epistasis in different environments. More than 90% of ncRNA interactions did not correlate with the epistasis of their neighbouring protein-coding genes, showing that the ncRNA and protein interaction networks are largely independent. In the presence of different stressors, epistatic changes were observed with an overall increase in negative interactions. Such network plasticity supports the notion that these ncRNAs may have environmentally dependent functions and can contribute to phenotypic adaptation. The U3 paralogs, SNR17A and SNR17B, share most interactions in rich media, as expected, however, in stressful conditions, there was an increase of paralog-unique interactions, suggesting that SNR17A and SNR17B may have diverged cellular roles under different environmental pressures and may have neo-functionalised after genome duplication. Further investigation of the interaction networks between SNR17 and protein-coding genes revealed the emergence of paralog-specific interactions under stress conditions. Gene Ontology enrichment analysis suggested a novel potential role of SNR17B in chromatin structural remodelling and DNA topology under osmotic and oxidative stress. Overall, this genome-wide study uncovers the level of epistasis involving ncRNA mutants aiding functional assignment and expanding our fundamental knowledge of these genetic elements.

## INTRODUCTION

The majority of biological processes in the cell are performed by proteins, either independently or through interactions with other proteins. However, eukaryotic genomes are pervasively transcribed, producing a wide range of non-coding RNAs (ncRNAs) that, although they are not translated into proteins, play important and often essential roles in cellular function (Chen & Kim, 2024; Jensen et al., 2013; Mattick et al., 2023; Qi et al., 2025; Wery et al., 2016; Wu et al., 2012). Approximately 98% of the human genome consists of non-protein-coding DNA sequences that were once regarded as non-functional evolutionary leftovers (Boland, 2017). In fact, most of the genome is transcribed and a number of ncRNAs are regulatory, assisting in essential cellular processes such as regulation of gene expression and transcription, apoptosis, telomere maintenance or RNA processing (Boland, 2017; Jensen et al., 2013; Qi et al., 2025; Wery et al., 2016; Wu et al., 2012). Since ncRNAs are integral components of cellular regulatory networks, it is not surprising that many diseases in humans, ranging from cancer to neurological disorders, are affected by mutations or defects in many of these ncRNAs (de Almeida et al., 2016).

In *Saccharomyces cerevisiae*, approximately 85% of the genome is transcribed, producing ca. 25% non-coding and ca. 75% protein-coding transcripts (Nagalakshmi et al., 2008). Due to the loss of the RNA interference (RNAi) machinery, *S. cerevisiae* does not possess short ncRNAs discovered in other eukaryotes, such as microRNAs (miRNAs) and small interfering RNAs (siRNAs) (Drinnenberg et al., 2009; Shabalina & Koonin, 2008). However, many classes of structured, regulatory and housekeeping ncRNAs are indeed present in the *S. cerevisiae* genome. These include tRNAs and rRNAs involved in protein synthesis, as well as small nuclear RNAs (snRNAs) known to perform intron splicing and small nucleolar RNAs (snoRNAs) that process or direct chemical modifications of other RNAs. Novel functions for these families of ncRNAs are still emerging. Furthermore, a great number of ncRNAs with unknown or unclear functions, so-called ‘non-conventional’ ncRNAs, have been discovered and classified into various groups depending on their half-life after transcription. The majority of ncRNAs are unstable (short-lived) as they are rapidly degraded by the RNA decay pathways. These include Cryptic Unstable Transcripts (CUTs) that were discovered in mutants of the nuclear exosome exonuclease component *RRP6* (Δ*rrp6*) (Neil et al., 2009; Xu et al., 2009), Xrn1 exo-ribonuclease sensitive Unstable Transcripts (XUTs) (van Dijk et al., 2011; Wery et al., 2016) and RNA-binding factor Nrd1 Unterminated Transcripts (NUTs) (Schulz et al., 2013). In contrast, there are also long-lived Stable Unannotated Transcripts (SUTs) that can evade nuclear degradation and are exported to the cytoplasm, where they are processed similarly to mRNAs and are present in wild-type cells (Xu et al., 2009). The stability of SUTs implies that they are functional and involved in cellular processes. It is estimated that in the genome of *S. cerevisiae* there are > 800 SUTs, > 900 CUTs, > 1800 XUTs and > 1500 NUTs (Schulz et al., 2013; Wery et al., 2016; Xu et al., 2009). These ncRNAs can be transcribed with the bidirectional promoter from both sense and antisense strands in relation to the coding genes from intra- and intergenic or open reading frame overlapping regions (Xu et al., 2009). Many of these ncRNAs overlap with each other and some are extended isoforms of other ncRNAs (Marquardt et al., 2011; Schulz et al., 2013; van Dijk et al., 2011; Wery et al., 2016).

In the past decade, large-scale studies to identify the cellular functions ascribed to ncRNAs have been performed (Balarezo-Cisneros et al., 2021; Huber et al., 2016; Parker et al., 2018; Schulz et al., 2013). For instance, libraries of 1502 barcoded ncRNA deletion mutants in *S. cerevisiae* in both haploid (*MAT**a*** and *MATα*) and diploid (homozygote and heterozygote) backgrounds were constructed as tools for ncRNA functional analysis (Parker et al., 2017, 2018). These libraries covered a total number of 443 unique ncRNAs deletion mutants, including tRNAs, snRNAs, snoRNAs, CUTs, SUTs and other annotated ncRNAs, that are localised to intergenic regions and do not overlap with protein-coding genes, their promoters or terminators.

While these resources enable systematic interrogation of ncRNA functions, single-mutant analyses alone provide limited insight into the functional relationships and genetic dependencies among ncRNAs. Large-scale genetic interaction mapping offers a complementary approach to uncover such relationships. This strategy has been extensively applied to protein-coding genes, enabling systematic quantification of epistasis across the genome (Baryshnikova et al., 2010; Tong & Boone, 2006). Systematic mapping of genetic interaction was first automated in *S. cerevisiae* by employing the single open reading frame deletion collection composed of ca. 4,800 genes crossed with 132 query strains using the Synthetic Genetic Array (SGA) methodology (Tong et al., 2004). To date, more than 23 million double mutants have been constructed and about 900,000 genetic interactions have been identified between thousands of protein-coding genes, revealing a large-scale genetic interaction map of the cell (Costanzo et al., 2016). Therefore, SGA analysis, which examines genetic interactions with ncRNAs, could serve as an efficient approach to explore ncRNA functions. Beyond direct ncRNA-associated interactions identified by SGA, deletion of a ncRNA may dysregulate its target protein-coding gene, and the combined expression changes of two such genes can produce a phenotype that is absent when either gene is transcriptionally altered alone. For example, this type of epistasis is evident upon SUT259/691 deletion, where *EMP46* and *GAL4* are both overexpressed, causing a lethal phenotype (Parker et al., 2018). Such epistatic interactions between proteins would not have been revealed by classical SGA analysis based on gene deletions.

Here, we employed the SGA-based approach to investigate the functions of ncRNAs in *S. cerevisiae* by constructing double ncRNA deletion mutants and analysing their genetic interactions. A mutant library comprising 379 ncRNAs was generated in the Y7092 query strain, and a subset of 34 deletions was crossed with 376 previously constructed ncRNA mutants (Parker et al., 2017, 2018). The resulting double mutants were assessed in rich media and four stress conditions (non-fermentable, oxidative, heat, and osmotic). A total number of 1,405 genetic interactions were identified in rich media, with ca. 42.9% displaying synthetic sick or lethal phenotypes. This predominance of positive interactions contrasts with protein-coding SGA networks, which are typically dominated by negative genetic interactions. These ncRNA interactions did not correlate with those of neighbouring protein-coding genes. We further identified 130 “core” interactions conserved across all conditions and a large set of environment-specific interactions, including distinct networks of the U3 snoRNA paralogs SNR17A and SNR17B, suggesting functional divergence driven by environmental adaptation. Additional SGA screening of both SNR17A and SNR17B mutants with a subset of the protein deletion collection (the diagnostic array) revealed the emergence of paralog-specific interactions under stress conditions. Further Gene Ontology enrichment analysis suggested a potential role of SNR17B in chromatin structural remodelling and DNA topology under osmotic and oxidative stress. This data supports the notion that SNR17A and SNR17B have neo-functionalized after the genome duplication event.

## RESULTS AND DISCUSSION

### Query ncRNA deletion mutant library construction and strain selection for SGA analysis

Single ncRNA deletion mutants in the Y7092 background (referred to here as ‘query’) were generated by replacing the ncRNA of interest with the *natMX4* marker that confers nourseothricin resistance (clonNAT) (see Supplementary Materials and Methods). A total of 379 deletion mutant query strains were generated. SGA analysis was conducted with 34 query strains (Supplementary Dataset S1), including 17 snoRNAs, 4 tRNAs and 13 SUTs and CUTs. These ncRNAs were chosen based on previous fitness studies that scored both the biomass as solid fitness of haploid ncRNA mutants, and the growth changes over time as a competitive fitness of diploid heterozygous ncRNA deletion mutants (Balarezo-Cisneros et al., 2021; Parker et al., 2018). Specifically, the deletion strains selected displayed either growth deficiency or haploinsufficiency under rich media or stressful conditions (*i.e.* high temperature, osmotic stress conditions, alternative carbon sources, high concentration of ethanol). According to previous work on protein-coding gene interaction maps, single mutant query strains with fitness defects are more likely to show a higher number of epistatic interactions compared to query strains with low or no fitness deficiency (Costanzo et al., 2010). In fact, there is a strong positive correlation between the number of genetic interactions and single-mutant fitness (Costanzo et al., 2010).

### SGA analysis of ncRNA mutants reveals a prevalence of positive epistasis in the genetic interaction network

We performed SGA screens by crossing the 34 selected ncRNA query strains with the ncRNA array library composed of 376 ncRNA deletion mutants (Supplementary Dataset S2), including 44 CUTs, 92 SUTs, 58 snoRNAs, 181 tRNAs and one unknown ncRNA RUF22. The SGA screen was repeated three times with three biological replicas (*i.e.* three independent transformants) of each query strain allowing the identification of reproducible epistatic interactions. Within each SGA screen, we had four to eight technical replicates of the array mutants and of the double mutants (see Materials and Methods). Out of ca. 13,000 interactions 37 were removed (ca. 0.3%) due to their inconsistent phenotypic behaviour between either biological or technical replicates. The linkage groups of ncRNAs that were on the same chromosome were also removed. For example, the snoRNAs SNR72, SNR73, SNR74, SNR75, SNR76, SNR77 and SNR78 are in a tight cluster on chromosome 13 and in the SGA were detected as synthetic lethal interactions because no recombination was occurring between them (spurious inter-cluster interactions). However, we also carried out SGA using the null mutant for the entire SNR72-78 cluster.

Approximately 1,405 out of the 12,516 double mutant combinations generated (ca. 11.2% of total interactions; Supplementary Dataset S3), displayed significant epistatic interactions (|ε| ≥ 0.15 and q ≤ 0.001; Figure 1, panel A). The query ncRNA had a median of 37.5 significant interactions, with numbers varying between 18 (for query SUT053-SUT468-CUT494) to 78 (for query SNR87) (Table 1 and Supplementary Dataset S3).

**Figure 1.**
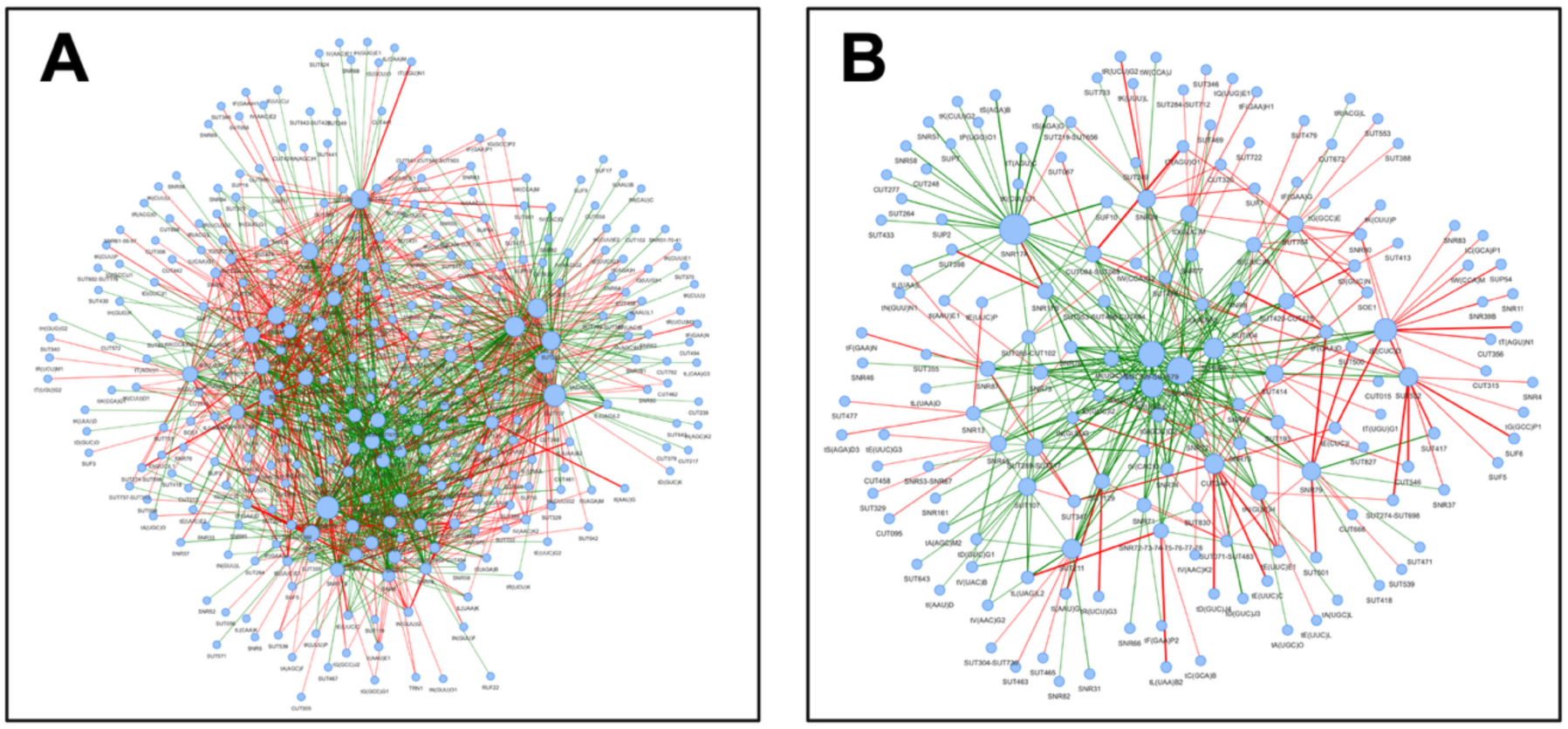
ncRNA SGA networks for 34 query strains. The SGA networks were generated based on a cut-off of (**A**) |ε| ≥ 0.15, q-value ≤ 0.001 and (**B**) |ε| ≥ 0.3, q-value ≤ 0.001 for 34 query ncRNA deletion strains. The node size is proportional to the number of interactions they are involved in. Red and green edges represent negative and positive epistasis, respectively. The thickness of the edges is proportional to the strength of the interaction.

**Table 1.** Number of significant SGA interactions between query and array strains, divided per ncRNA classes. Positive and negative epistasis are represented with + and -, respectively. Refer to Supplementary Dataset S3 for detailed interactions.

| Query | Number of interactions in the different ncRNA classes |  |  |  |  |  |  |  | Total interactions |  |  |
| --- | --- | --- | --- | --- | --- | --- | --- | --- | --- | --- | --- |
|  | CUTs |  | SUTs |  | SNRs |  | tRNAs |  | + | - | + & - |
|  | + | - | + | - | + | - | + | - |  |  |  |
| CUT084-SUT068 | 0 | 4 | 4 | 4 | 2 | 3 | 16 | 7 | 22 | 18 | 40 |
| CUT244 | 0 | 1 | 4 | 5 | 2 | 6 | 8 | 8 | 14 | 20 | 34 |
| SNR13 | 1 | 2 | 5 | 4 | 3 | 4 | 9 | 3 | 18 | 13 | 31 |
| SNR17A | 3 | 1 | 7 | 11 | 6 | 8 | 22 | 4 | 38 | 24 | 62 |
| SNR17B | 0 | 1 | 5 | 3 | 2 | 1 | 12 | 7 | 19 | 12 | 31 |
| SNR34 | 2 | 3 | 5 | 2 | 5 | 5 | 15 | 11 | 27 | 21 | 48 |
| SNR45 | 3 | 7 | 8 | 9 | 6 | 3 | 14 | 15 | 31 | 34 | 65 |
| SNR71 | 1 | 1 | 6 | 5 | 3 | 0 | 11 | 4 | 21 | 10 | 31 |
| SNR72 | 1 | 0 | 5 | 5 | 0 | 0 | 7 | 12 | 13 | 17 | 30 |
| SNR72-73-74-75-76-77-78 | 0 | 2 | 10 | 2 | 5 | 1 | 12 | 9 | 27 | 14 | 41 |
| SNR73 | 0 | 2 | 6 | 4 | 2 | 0 | 16 | 5 | 24 | 11 | 35 |
| SNR74 | 0 | 0 | 9 | 3 | 3 | 2 | 10 | 6 | 22 | 11 | 33 |
| SNR75 | 2 | 0 | 5 | 6 | 3 | 0 | 12 | 9 | 22 | 15 | 37 |
| SNR76 | 1 | 1 | 9 | 4 | 2 | 0 | 9 | 4 | 21 | 9 | 30 |
| SNR77 | 0 | 0 | 11 | 5 | 1 | 0 | 12 | 7 | 24 | 12 | 36 |
| SNR78 | 1 | 0 | 6 | 3 | 1 | 0 | 10 | 3 | 18 | 6 | 24 |
| SNR79 | 1 | 0 | 12 | 6 | 4 | 0 | 12 | 8 | 29 | 14 | 43 |
| SNR8 | 0 | 0 | 8 | 5 | 1 | 0 | 12 | 6 | 21 | 11 | 32 |
| SNR87 | 2 | 3 | 10 | 9 | 11 | 1 | 19 | 23 | 42 | 36 | 78 |
| SUT053-SUT468-CUT494 | 0 | 0 | 5 | 3 | 1 | 0 | 7 | 2 | 13 | 5 | 18 |
| SUT107 | 7 | 6 | 12 | 9 | 7 | 1 | 18 | 13 | 44 | 29 | 73 |
| SUT129 | 0 | 0 | 5 | 3 | 1 | 0 | 9 | 8 | 15 | 11 | 26 |
| SUT193 | 1 | 6 | 9 | 1 | 2 | 2 | 7 | 8 | 19 | 17 | 36 |
| SUT211 | 4 | 4 | 7 | 9 | 9 | 1 | 18 | 8 | 38 | 22 | 60 |
| SUT289-SUT717 | 3 | 4 | 10 | 7 | 12 | 1 | 16 | 18 | 41 | 30 | 71 |
| SUT414 | 1 | 5 | 8 | 7 | 2 | 2 | 9 | 7 | 20 | 21 | 41 |
| SUT420-CUT425 | 0 | 1 | 8 | 9 | 3 | 5 | 10 | 7 | 21 | 22 | 43 |
| SUT496 | 1 | 4 | 8 | 2 | 1 | 2 | 6 | 2 | 16 | 10 | 26 |
| SUT532 | 1 | 5 | 6 | 2 | 2 | 5 | 12 | 9 | 21 | 21 | 42 |
| SUT764 | 0 | 2 | 9 | 7 | 2 | 3 | 9 | 8 | 20 | 20 | 40 |
| tD(GUC)M | 0 | 3 | 5 | 11 | 4 | 2 | 13 | 7 | 22 | 23 | 45 |
| tE(CUC)D | 0 | 3 | 6 | 7 | 2 | 7 | 12 | 11 | 20 | 28 | 48 |
| tE(UUC)M | 1 | 3 | 6 | 7 | 6 | 1 | 8 | 6 | 21 | 17 | 38 |
| tH(GUG)H | 0 | 1 | 7 | 6 | 1 | 4 | 10 | 8 | 18 | 19 | 37 |
| **Total** | **37** | **75** | **246** | **185** | **117** | **70** | **402** | **273** | **802** | **603** | **1,405** |

In contrast to protein-coding genes, for which negative genetic interactions are approximately two-fold more prevalent than positive interactions (Costanzo et al., 2010), here there were more double mutants displayed positive epistasis (ca. 57.1%; Table 1) than negative (Table 1). This trend is exacerbated if the stringency of the epsilon is doubled to 0.3, with 64.7% (246 out of 380) of the interaction being positive (Figure 1, panel B).

Cases of synthetic lethality were also identified (Supplementary Dataset S3). The paralogs SNR17A and SNR17B, the U3 snoRNAs responsible for 35S pre-rRNA cleavage and 18S rRNA maturation, were found to be synthetic lethal as expected (Hughes et al., 1987). The tRNAs, tE(CUC)I and tE(CUC)D, both members of the same two-copy tRNA family, were also found as expected to be synthetically lethal due to the complete loss of the tRNA family function (Bloom-Ackermann et al., 2014). CUT244 exhibited synthetic lethality with tE(UUC)E1 and tE(UUC)E2, both belonging to the wobble uridine tRNA family that decodes glutamate. SNR79 on chromosome XII displayed synthetic lethality with two neighbouring ncRNAs, SUT500 and SUT501, on chromosome V suggesting a functional association with the corresponding chromatin region on this chromosome.

When looking at the classes of ncRNAs in the array, we found a higher frequency of SUTs involved in the interactions than CUTs, with an average of 4.5 and 2.3 interactions per ncRNA, respectively. This observation is consistent with SUTs being transported to the cytoplasm, similarly to mRNAs, and acting as functional transcripts affecting important cellular functions (Balarezo-Cisneros et al., 2021; Kyriakou et al., 2016; Tuck & Tollervey, 2013), therefore placing them more centrally within genetic networks than other classes of ncRNAs.

### Comparison between ncRNA and neighbouring protein interaction networks reveals poor overlap

To understand whether a potential alteration of expression of a gene flanking a ncRNA deletion may have indirectly influenced the ncRNA interaction network, we surveyed all genetic and physical interactions of the proteins encoded by the neighbouring genes, to determine the overlap between protein and ncRNA interaction networks. Intergenic ncRNAs could either work in *trans* affecting genes or transcription factors involved in cellular growth (Balarezo-Cisneros et al., 2021; Camblong et al., 2009; Parker et al., 2018) or may provoke expressional changes in *cis* (Camblong et al., 2007; Martens et al., 2004; Thebault et al., 2011; van Werven et al., 2012) with this expression change potentially causing the epistasis between the affected genes (*i.e.* synthetic interactions between the neighbouring genes due to expressional changes). Provided that the latter scenario is the most common (Qi et al., 2025), we would expect an overlap between the SGA network of ncRNAs and the SGA network of the neighbouring protein-coding genes.

The BioGrid database was used to recover the interactions which are recorded as genetic or physical (*i.e.* protein-protein interactions scored either by affinity capture, protein-fragment complementation assay or two-hybrid bait). We found that the overwhelming majority of neighbouring genes (94.9%), which are flanking the genetically interacting ncRNAs, had no recorded genetic or protein interactions (Figure 2 and Supplementary Dataset S4). The proportion of ncRNAs interacting pairs which are flanked by interacting proteins was ca. 5.1 %. This percentage is ca. 20% lower than the probability that any of the four neighbouring protein-coding genes of two randomly selected ncRNAs would interact by chance (null expectation of 6.32%). These data suggest that the ncRNA genetic interaction network is not shaped solely by the *cis* interactions between ncRNAs and their neighbouring coding genes, and that *trans* effects may be more prevalent than suggested by current models of ncRNA function (Qi et al., 2025).

**Figure 2.**
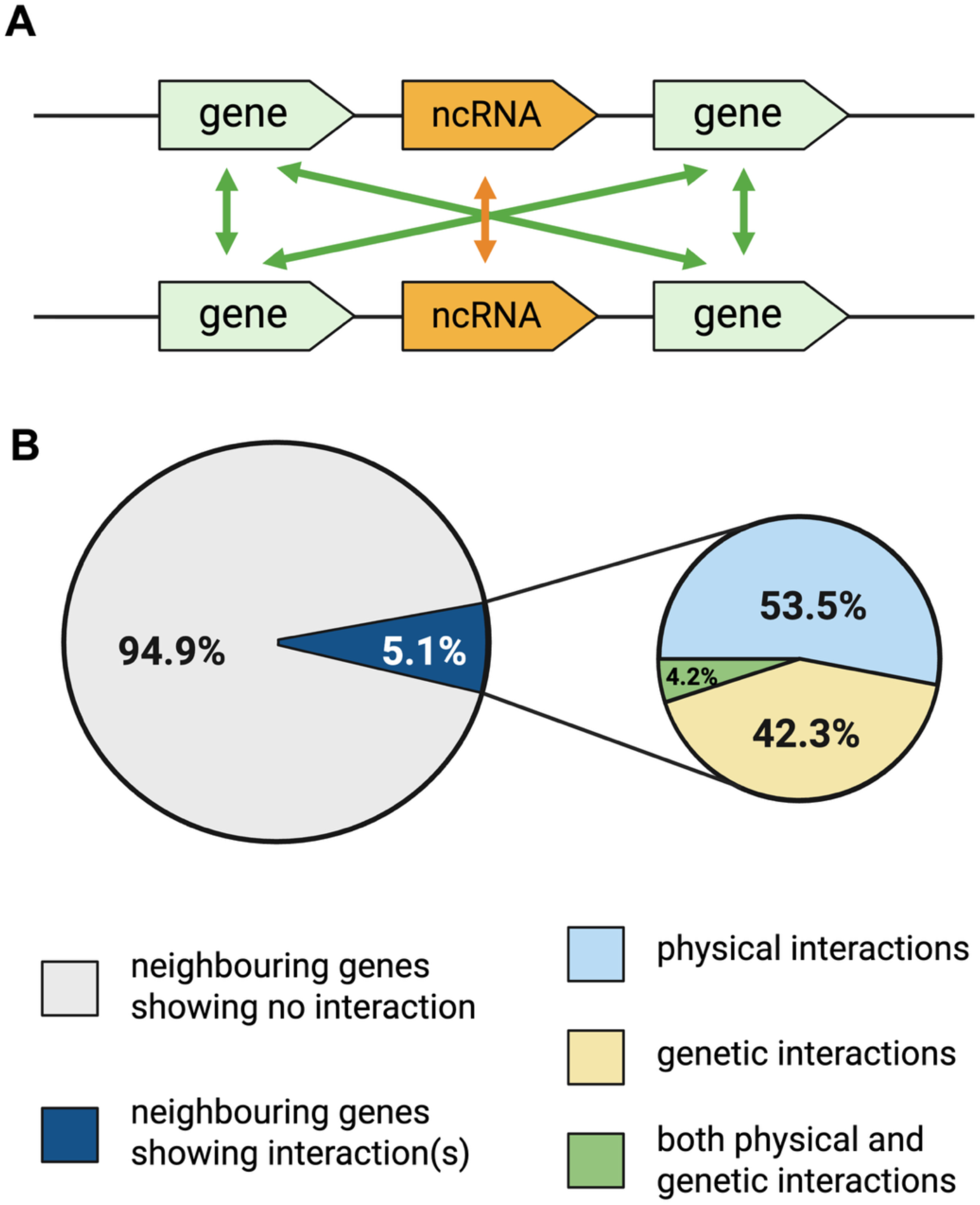
Analysis of the interactions of genes neighbouring epistatic pairs of ncRNAs. (**A**) Illustration of the analytical workflow used to investigate genetic and physical interactions between the neighbouring protein-coding genes (green arrows) of ncRNA pairs showing significant epistasis (orange). (**B**) Pie charts displaying the proportion of significantly interacting neighbouring genes (dark blue colour), broken down in genetic (light yellow), physical interactions (light blue) or with both interaction (green).

Indeed, recent data from transcriptome studies of ncRNA mutants revealed that the deletion of several SUTs has a global, as well as a local effect on gene transcription, and can act on a large-scale by modulating the expression of transcription factors (TFs) (Balarezo-Cisneros et al., 2021). For instance, in the absence of SUT532, SUT125 and SUT126, the expression of TFs that participate in the stress response, sporulation, cell cycle progression and cell integrity, such as *XPB1, RIM1 RGM1, YOX1, ROF1, TOS8, MSA1,* and *TOS4* was highly affected. Such modulation is scored as a growth advantage in the short term, although unlikely to render the cell fitter over time given that an accurate mitotic control is crucial to avoid multiple duplications and DNA damage (Balarezo-Cisneros et al., 2021).

The subset of neighbouring gene pairs associated with interacting ncRNAs, suggests a potential *in cis* effect of the ncRNA on the flanking genes (*i.e.* ncRNAs that are affecting the regulation of local genes). When generating the ncRNA mutant collection (Parker et al., 2017, 2018) every effort has been made to avoid deletion of either promoter or terminators of the adjacent protein-coding genes, but it is possible that for some ncRNA double mutants the observed phenotype is due (or partially due) to inadvertent partial deletions of other regulatory elements.

Interestingly, in the subset of potentially *cis*-acting ncRNAs, we detected an enrichment of positive interactions when the neighbouring genes exhibit physical interactions (25 of 33, ca. 75.7%; Supplementary Dataset S4). This data supports the previous observation that physically interacting proteins are often associated with positive genetic interactions (Costanzo et al., 2010).

### Network dynamics of ncRNAs in different stress conditions exhibit more negative epistatic interactions

Genetic epistasis is dependent on the genetic background and the environment (van Leeuwen et al., 2016). Therefore, we analysed all single and double ncRNA mutants under five different nutritional and stress conditions (rich and non-fermentative media as well as oxidative, high temperature, and osmotic stresses) to detect environmental-specific deviation of growth and infer changes in the ncRNA network. We scored the fitness plasticity of the interaction networks with 29 query strains encompassing ca. 11,000 mutant combinations (Table 2 and Supplementary Dataset S5). At this stage, some haploid array single mutants failed to grow on glycerol, indicating mitochondrial DNA loss during storage or culturing. These single mutants, along with their corresponding double mutants, were removed from the entire dataset, as mitochondrial deficiency strongly affects haploid phenotypes and biases epistasis toward positive when crossed with the query strain.

**Table 2:** Number of significant ncRNA interactions after fitness profiling in different environmental conditions.

| Condition | Total number of significant interactions | Number of positive interactions | Number of negative interactions |
| --- | --- | --- | --- |
| YPD, 30 °C | 1,067 | 579 | 488 |
| YPD, 37 °C | 761 | 349 | 412 |
| YP + 7% glycerol, 30 °C | 4,412 | 2,090 | 2,322 |
| YPD + 1 M NaCl, 30 °C | 1,037 | 495 | 542 |
| YPD + 4 mM H<sub>2</sub>O<sub>2</sub>, 30 °C | 1,153 | 506 | 647 |

$|\epsilon| \geq 0.15$ , q-value $\leq 0.001$

The stress conditions produce an overall increase in the proportion of negative interactions (ca. 52.2%–56.1% according to different conditions) compared to YPD 30 °C (ca. 45.7%, Supplementary Figure S1). This negative interaction trend was consistent when we raised the stringency of the study adapting the value of 0.3 as an absolute epsilon threshold (Figure 3, Supplementary Dataset S5). In this case, ca. 63.3%–84.6% negative interactions were observed under stress conditions, compared with 32.9% in rich media. The highest number of total interactions was observed under respiratory stress (YP + 7% glycerol, 30 °C; Figure 3 panel C), where there was ca. 4 times more negative interactions than in rich medium.

**Figure 3.**
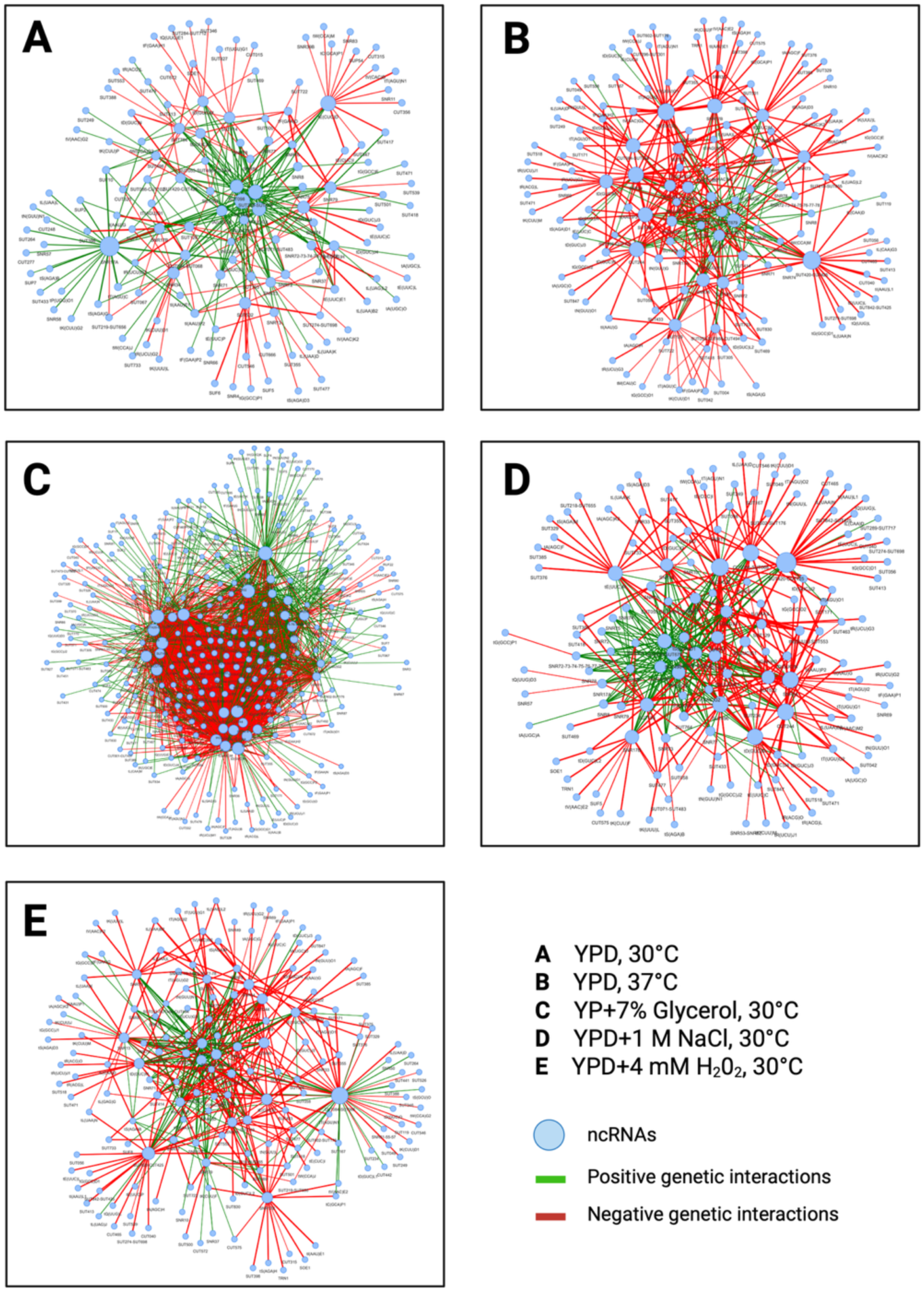
Visualisation of the ncRNA network under different environmental conditions. ncRNA network generated for 29 query ncRNA deletion strains based on the *|ε|* ≥ 0.3 and q-value ≤ 0.001 cut off in (**A**) YPD, 30 °C, (**B**) YPD, 37 °C, (**C**) YP + 7% Glycerol, 30 °C, (**D**) YPD + 1 M NaCl, 30 °C and (**E**) YPD + 4 mM H_2_O_2_, 30 °C. Network ncRNAs nodes (blue); negative ncRNA genetic interactions (red edges); positive ncRNA genetic interactions (green edges). The thickness of the edges is proportional to the absolute epsilon.

Next, we identified ncRNA interactions that were shared and unique to each of the environmental conditions tested (|ε| ≥ 0.15 and q-value ≤ 0.001) (Figure 4, panel A, Supplementary Dataset S6). Three hundred and seventy-two ncRNA interactions were specific only to rich medium (YPD, 30 °C), 2,971 to glycerol (respiratory stress), 203 to NaCl (osmotic stress), 45 to 37 °C (temperature stress) and 266 to hydrogen peroxide (oxidative stress).

**Figure 4.**
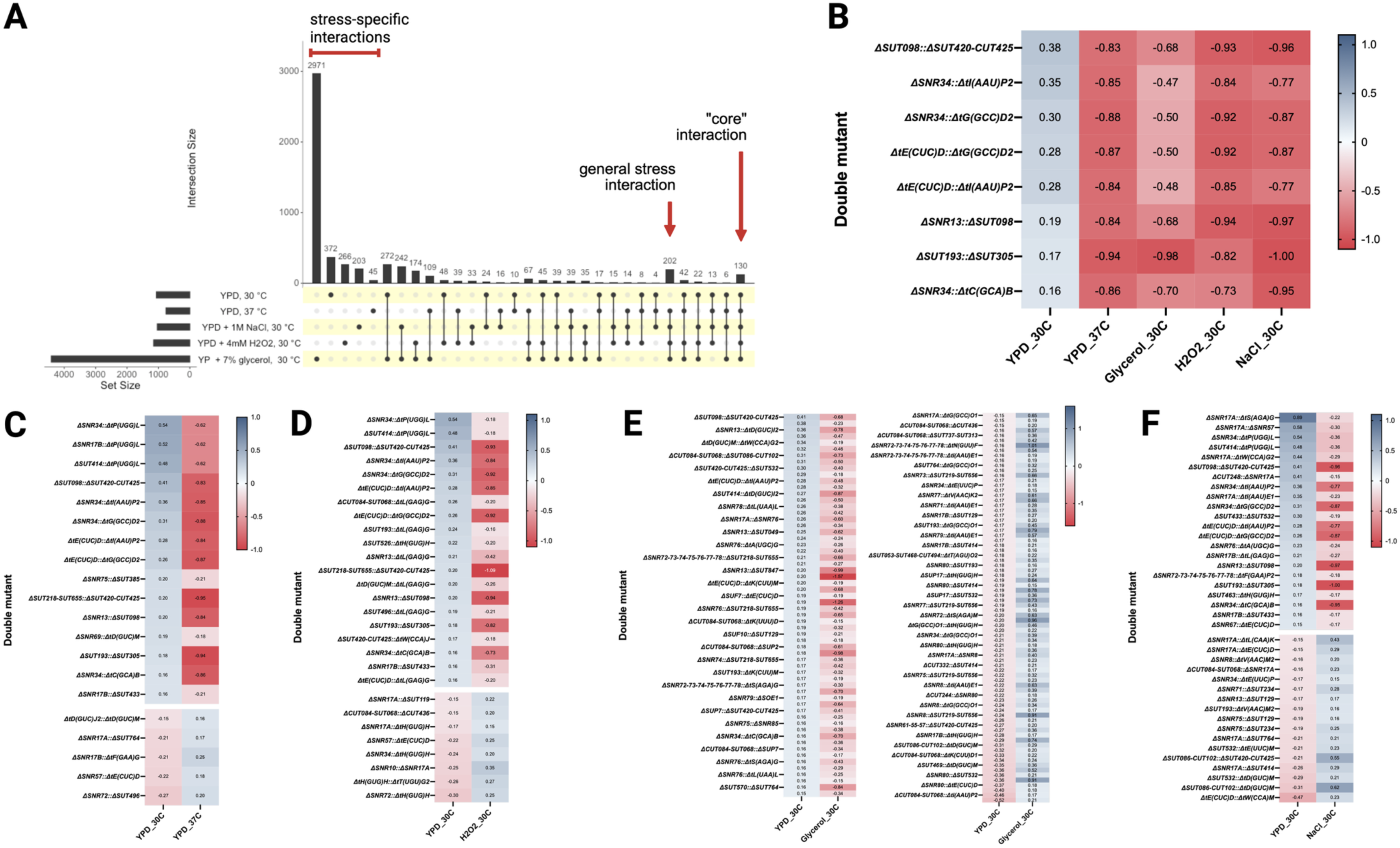
Environmental conditions elicit yeast ncRNA interaction network changes. (**A**) Upset plot depicting environmental unique and shared ncRNA genetic interactions between the five conditions tested (YPD, 37 °C; YPD, 30 °C; YPD + 1 M NaCl, 30 °C; YPD + 4 mM H_2_O_2_, 30 °C; and YP + 7% glycerol, 30 °C). The bottom left-hand side horizontal bar chart displays the total number of significant interactions for each condition tested. The top vertical bar chart displays the number of interactions unique or shared between conditions, indicated by a black dot in the bottom chart box. Unique interactions are represented by a single black dot between all 5 conditions tested. Several dots connected with a black line represent the number of interactions shared in the selected conditions. (**B**) Heatmap depicting epsilon of eight ncRNA interactions that display significant change in the four stress conditions tested compared to the rich medium. Specific interactions are arranged on the y-axis. Conditions are displayed on the x-axis. Negative interactions are represented as shades of red. Positive interactions are represented as shades of blue. **(C*–*F)** Heatmap depicting epsilon of ncRNA interactions that display significant change in between the standard condition (YPD at 30 °C) and one of the stress condition, including (**C**) YPD at 37 °C; (**D**) YPD + 4 mM H_2_O_2_ at 30 °C; (**E**) YP + 7% glycerol at 30 °C; and (**F**) YPD + 1 M NaCl at 30 °C.

We found 130 “core” ncRNA interactions that were shared between all five conditions tested (Supplementary Figure S2). These data allowed us to identify SUTs and CUTs (77 interactions, 59.2%) that may play a more central role in maintaining the ncRNA interaction network across the conditions. Interestingly, within the core ncRNAs interactions, there were eight that in the stress conditions changed from positive to negative compared to the rich medium (Figure 4, panel B). Several environment-specific interactions also exhibited direction changes compared with YPD 30 °C (Figure 4, panel C–F). More interactions shifted from positive to negative under high-temperature and oxidative stress, while under osmotic stress and glycerol conditions, direction changes were more balanced.

The sub-networks with shared common ncRNAs that change directionality in the stress conditions suggested that these ncRNAs may have a more prominent function in stress response. We identified a prevalence of negative interactions under obligatory respiratory conditions (YP + 7% glycerol, 30 °C) among ncRNAs that had previously been described to display a pivotal role in cell fitness (Balarezo-Cisneros et al., 2021). For example, SUT532 appeared to modulate genes *in trans* that are involved in mitochondrial functions and display 108 unique interactions with other ncRNAs in glycerol, of which 53 are negative interactions (Supplementary Dataset S6).

Additionally, we identified 202 shared interactions in common to all four stressed conditions but not in YPD at 30 °C (Figure 4, panel A). In this group, snoRNAs were under-represented (2.5% out of 13.3% present in the total interactions), and none of the 202 stress-specific interactions were observed between snoRNA pairs. Here, the low number of snoRNA interactions underlined the core cellular function of snoRNAs and revealed a less dynamic network across environmental conditions compared with those of other ncRNA classes.

We next carried out correlation studies between the number of genetic interactions for individual ncRNAs mutants and their phenotypes, using previously published fitness data (Balarezo-Cisneros et al., 2021). Unlike the protein network where the correlation between number of interactions and phenotype is strong in rich media at 30 °C, here we did not find significant correlation between these two parameters in both rich media (Supplementary Figure S3, panel A) or in stressful environments (Supplementary Figure S3, panel B,C,D). This lack of correlation suggests that the extent of genetic interactions cannot be simply explained by the fitness defect of single deletions but rather reflects additional functional dynamics of ncRNAs within the cellular network.

For each stress condition, we also compared the ncRNA interactions with the only available interaction network of the neighbouring protein-coding genes (in YPD at 30 °C), using the BioGRID database. Again, only a small subset of neighbouring genes/proteins displayed interactions either genetic or physical (Supplementary Figure S4 and Supplementary Dataset S7).

### Analysis of the snoRNAs paralogs, SNR17A and SNR17B

In *S. cerevisiae,* the SNR17 snoRNA is transcribed from two genetic loci, SNR17A and SNR17B, located on separate chromosomes, XV and XVI, respectively. SNR17A and SNR17B both encode the U3 snoRNA, which is required for pre-rRNA processing, namely cleavage of 35S pre-rRNA leading to mature 18S rRNA production (Maxwell & Fournier, 1995; Woolford & Baserga, 2013). An early study suggested that these two paralogs share the same nuclear localisation and bind to 35S pre-rRNA (Hughes et al., 1987) and hence may perform the same function in the cell. In fact, they can compensate, or partially compensate, each other’s mutation in rich medium. However, the double deletion of SNR17A and SNR17B is lethal (Hughes et al., 1987) and our SGA results confirm this outcome.

In YPD at 30 °C, and 37 °C the deletion of SNR17B does not impact the phenotype compared to the WT, while a small fitness defect is detected in the SNR17A deletion strain, suggesting only a partial compensation from the other paralog. A worse fitness cost for ΔSNR17A was observed under oxidative stress conditions, suggesting that in this case SNR17B cannot properly compensate for the SNR17A deletion (Supplementary Figure S5). In YP supplemented with 7% glycerol, both ΔSNR17A and ΔSNR17B exhibited comparable fitness defects relative to the WT, while in YPD with 1 M NaCl, no difference in fitness was detected with either snoRNA mutant versus the WT. The data supported the notion that the compensation function of the paralogs is environmental dependent.

The SGA network of ΔSNR17A and ΔSNR17B can help to understand whether the function of these two paralogs has diverged over evolutionary time. Seventy-one and 31 significant SGA interactions were detected in our screens in YPD at 30 °C with ΔSNR17A and ΔSNR17B respectively (Figure 5, panel A and Supplementary Dataset S8). The higher number of total interactions with SNR17A is consistent with the observation that single mutants with fitness defects tended to exhibit an increased number of interactions as reported in (Costanzo et al., 2010). As expected, a number of interactions were shared between these two snoRNAs, reflecting that they are functional homologs, and specifically SNR17B shared more than half of its interactions with SNR17A (Figure 5, panel A and F). The number of unique interactions for SNR17A was approximately three times higher than that for SNR17B, consistent with the observation that deletion of SNR17A causes a greater fitness reduction than deletion of SNR17B (Supplementary Figure S5). However, an increase in SNR17B-specific interactions was observed in other environmental conditions (Figure 5, panels B, C, D and E). The expansion of the SNR17B interaction network was also detected when the most stringent epsilon value of 0.3 was used (in the main text Figure 5, panels G, H, I and J; and Supplementary Dataset S8). This difference in the interaction network between SNR17A and SNR17B may indicate that, since the duplication event, the paralogs may have neo-functionalised, and SNR17B might also be playing a role in other stress-induced functions independently of its paralog SNR17A.

**Figure 5.**
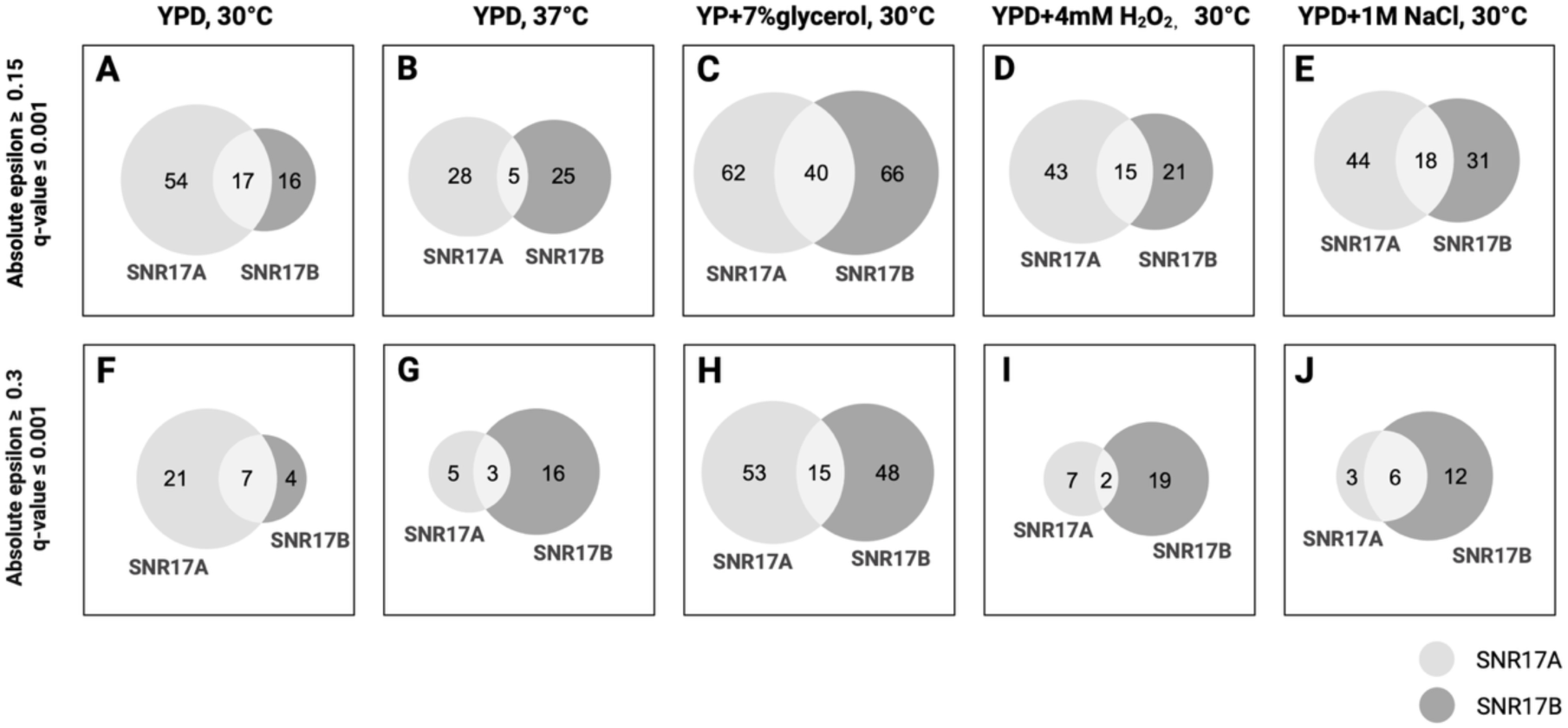
Area proportional Venn diagrams displaying unique and shared interactions between SNR17A and SNR17B in ncRNA SGA. Unique and shared interactions between SNR17A and SNR17B are displayed in (A & F) YPD, 30 °C; (B & G) YPD, 37 °C; (C & H) YP + 7% glycerol, 30 °C; (D & I) YPD + 4 mM H_2_O_2_, 30 °C and (E & J) YPD + 1 M NaCl, 30 °C. Light grey (centre) represented shared interactions between SNR17A and SNR17B; mid grey (left) represented unique interactions to SNR17A; dark grey (right) represented unique interactions to SNR17B.

### SGA analysis of U3 snoRNA with protein-coding genes reveals an overlap of genetic interactions of SNR17A and SNR17B in rich media

To further investigate the interaction network of the U3 snoRNA paralogs, we performed SGA analysis with the query strains ΔSNR17A and ΔSNR17B and a diagnostic collection of protein-coding genes, which includes 1032 non-essential gene mutants representing different cellular pathways (Kuzmin et al., 2021). Because temperature-sensitive (ts) mutants were present in the collection, the experiments were performed at the permissive temperature of 26 °C (rather than 30 °C). The double mutants were created using 3 biological replicates for each query (*i.e.* three independent transformants), resulting in 2,462 double mutant combinations per biological replica.

Similar to the SGA analysis of ncRNA mutants, genes located within 2 kb of the SNR17A or SNR17B loci were excluded, including *HES1* for SNR17A, and *KES1* and *POC4* for SNR17B. As expected, *HES1* did not form any viable double mutant with SNR17A mutant, whereas *KES1* and *POC4* failed to form viable double mutants with SNR17B mutant, which validates the robustness of our SGA. In addition, genes associated with the auxotrophic markers used during SGA, including *LYS1, LYS2, ARG4, ARG56, HIS1* and *HIS7*, were excluded from the dataset, as strains carrying these mutations cannot survive the SGA selection process.

There are 54 significant (absolute |ε| ≥ 0.15, q-value ≤ 0.001) genetic interactions observed for SNR17A (23 positive and 31 negative; Table 3, Supplementary Dataset S9) and 35 significant genetic interactions observed for SNR17B (13 positive and 22 negative interaction; Table 3 and Supplementary Dataset S9). Again, the larger number of interactions with SNR17A is consistent with the observation that single mutants with fitness defects tended to exhibit an increased number of interactions (Costanzo et al., 2010).

**Table 3.**
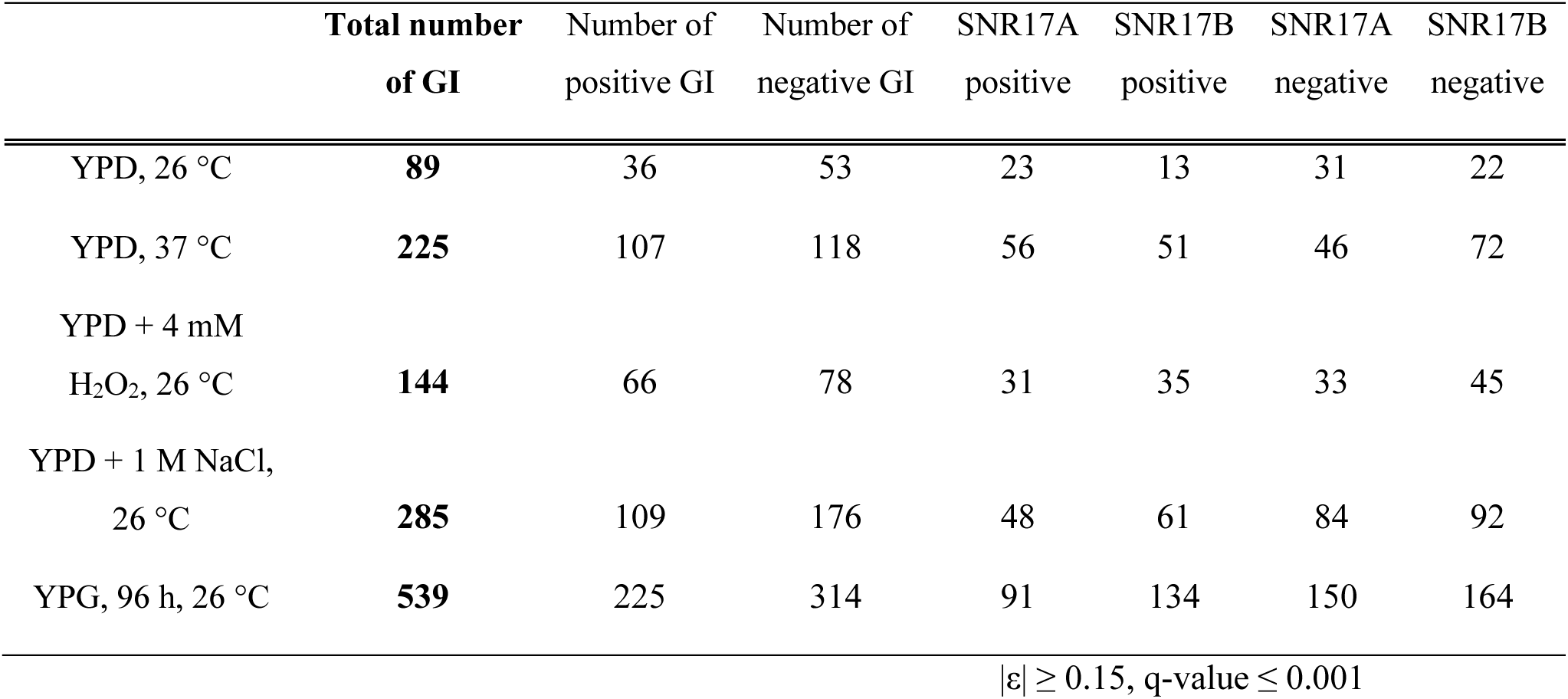
Number of significant genetic interactions (GI) in different environmental conditions.

In total, 18 interactions were shared between SNR17A and SNR17B, accounting for 33% of SNR17A interactions and 51% of SNR17B interactions (Figure 6, panel A). This trend persisted at the most stringent epsilon threshold of 0.3 (Figure 6, panel F). These observations support the notion that the function of SNR17B largely overlaps with that of SNR17A under rich media conditions, and that SNR17A is the primary paralog that carries out function.

**Figure 6.**
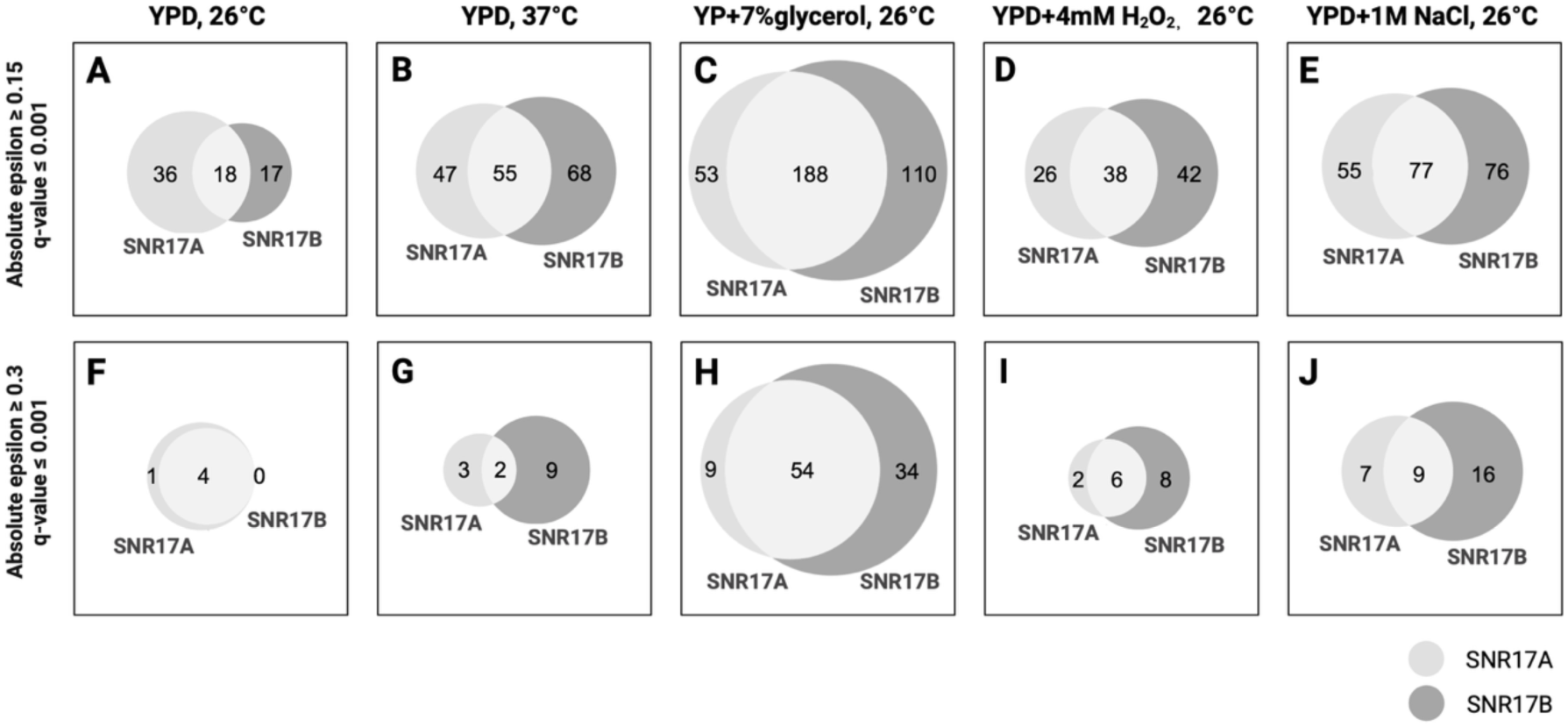
Area proportional Venn diagrams displaying unique and shared interactions between SNR17A and SNR17B in protein coding gene SGA. Unique and shared interactions between SNR17A and SNR17B (q ≤ 0.001) are displayed in (A & F) YPD, 26 °C; (B & G) YPD, 37 °C; (C & H) YP + 7% glycerol, 26 °C; (D & I) YPD + 4 mM H_2_O_2_, 26 °C and (E & J) YPD + 1 M NaCl, 26 °C. Light grey (centre) represented shared interactions between SNR17A and SNR17B; mid grey (left) represented unique interactions to SNR17A; dark grey (right) represented unique interactions to SNR17B.

We then mined the BioGRID database to pull out data of known interactions between U3 snoRNAs and coding genes. Among the 46 genes reported to have genetic or physical interactions with SNR17, two were present in our array, namely *RPS18B* and *HHF2*, which physically interact with U3 snoRNA. *RPS18B* encodes a protein component of the small (40S) ribosomal subunit (Lecompte et al., 2002) and *HHF2* encodes for the histone protein H4 (Megee et al., 1995). In our SGA, both SNR17A and SNR17B exhibited weak but significant positive interactions with *RPS18B* as expected (ε = 0.11 with SNR17A and ε = 0.06 with SNR17B, both q < 0.001). No interaction was detected between U3 and *HHF2*, reflecting the limited overlap between physical and genetic interaction networks. In fact, positive genetic interactions between nonessential genes only partially overlap with known protein-protein interactions (Costanzo et al., 2016).

We next examined genes exhibiting strong paralog-specific interactions, such as those detected only with SNR17A or only with SNR17B. Specifically, *SLX9*, a gene encoding a nuclear export receptor that is involved in the maturation of ribosomal 18S rRNA (Bax et al., 2006), only interacts with SNR17A, showing a strong positive interaction (ε = 0.31; Supplementary Dataset S9, Figure 7, panel A). Slx9p has been shown to physically associate with U3 snoRNA, specifically, ProtA–Slx9p co-precipitates substantial amounts of U3 snoRNA (Bax et al., 2006), indicating that Slx9p participates in a complex involving U3 during small subunit rRNA maturation. Given that no interaction between *SLX9* and SNR17B was observed (ε = 0.009, Figure 7, panel A), SNR17B may not be involved in the function associated with the rRNA export and processing pathway. Slx9p has been identified as a G-quadruplex (G4) binding protein that recognises guanine-rich secondary nucleic acid structures (Götz et al., 2019). G4 structures are formed at specific G-rich regions containing at least four guanine tracts, with each tract containing three or more consecutive guanines (G_≥3_N_x_G_≥3_NₓG_≥3_NₓG_≥3_) (Meier-Stephenson, 2022). Comparison of the sequences of the two paralogs revealed four guanine tracts located near the 3′ end of SNR17A, while only three guanine tracts were present in SNR17B (Figure 7, panel B, highlighted in red). This difference may account for the reduced association of SNR17B with Slx9p.

**Figure 7.**
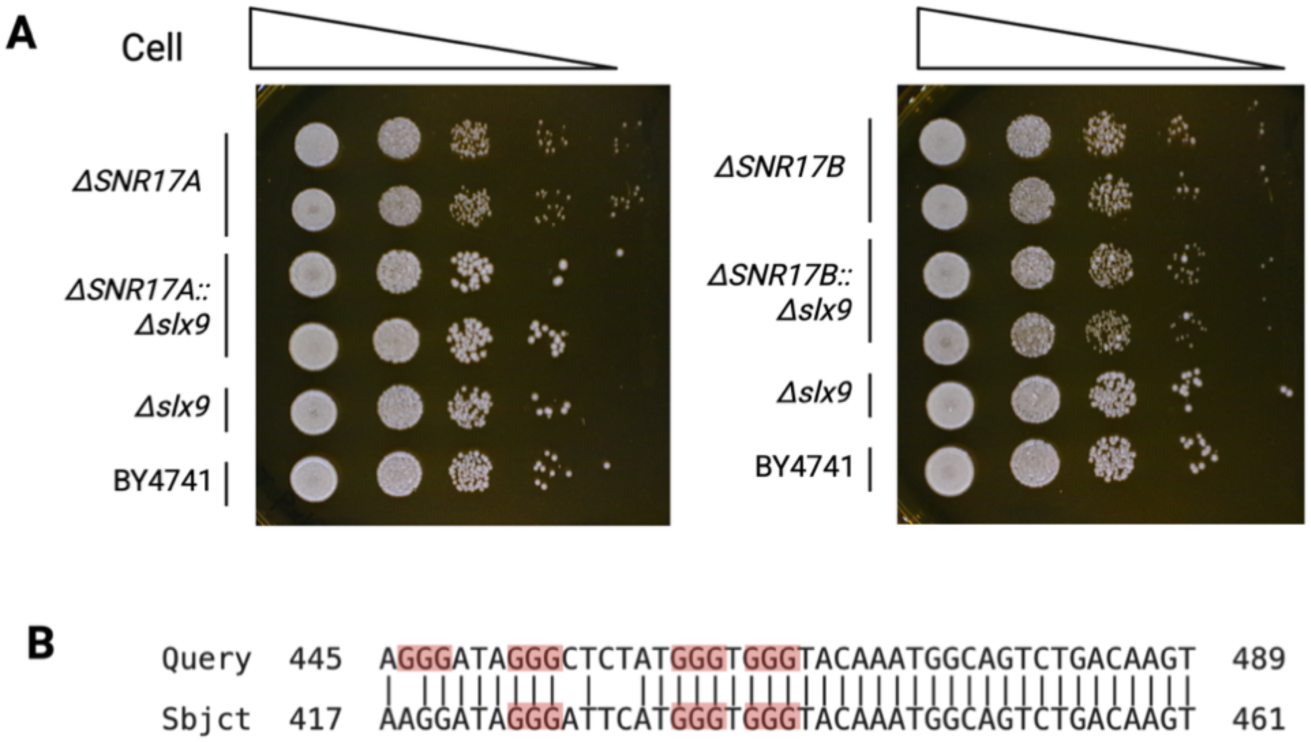
SNR17 paralog-specific interactions with *SLX9.* (**A**) validation for positive GI between SNR17A and *SLX9*. Spot assay of ΔSNR17A or ΔSNR17B and Δslx9, performed on YPD at 26°C, single mutant and double mutants. (**B**) sequence alignment of 3’ end of SNR17A and SNR17B. Guanine tracts (G≥3) are highlighted in red.

Among the shared top strong interactions between SNR17A and SNR17B, there are genes such as *VAM3* encoding SNAP receptor subunit of a vacuolar membrane protein SNARE complex (Wada et al., 1992); *AVL9* encoding for protein involved in post-Golgi vesicle-mediated transport (Harsay & Schekman, 2007); *YME1* encoding for catalytic subunit of i-AAA protease complex that is located in mitochondrial inner membrane; and *GEM1* encoding mitochondrial membrane GTPase (Frederick et al., 2004) (Supplementary Dataset S9, in bold). These genes are not directly related to U3 snoRNA function in rRNA processing and ribosome biogenesis, but are all involved in membrane-associated processes. Increasing evidence has suggested a functional coupling between ribosome synthesis and membrane-associated processes (B. Li & Warner, 1996; Mizuta & Warner, 1994), particularly along the secretory pathway, from peptide insertion into the endoplasmic reticulum (ER) to vesicle fusion with the plasma membrane (Y. Li et al., 2000). The presence of these interactions in both SNR17A and SNR17B networks may therefore reflect this connection between U3 snoRNA function and broader membrane-associated processes.

### SNR17 exhibited more genetic interactions in different stress conditions

Following the SGA analysis, we next examined the plasticity of the interaction networks under four additional stress conditions. Similar to the ncRNA-ncRNA SGA interaction network, an overall increase in the total number of interactions was observed under stress, especially for SNR17B (Table 3; Figure 6, panel B–E). Similar expansion of the SNR17B interaction network was also observed at the epsilon threshold of 0.3 (Figure 6, panel G–J). Among the four stress conditions, the largest increase was detected in YP with 7% glycerol, where the number of interactions was approximately six-fold higher than in rich media. In this YP with 7% glycerol condition, 142 interactions were identified for SNR17A and 175 interactions for SNR17B (Figure 8, panel A). We also examine the direction of genetic interaction in each stress condition. No significant change was observed in the overall ratio of positive to negative interactions for the total of SNR17A and SNR17B interactions under stress conditions (two-proportion Z-test, p-value = 0.87).

**Figure 8.**
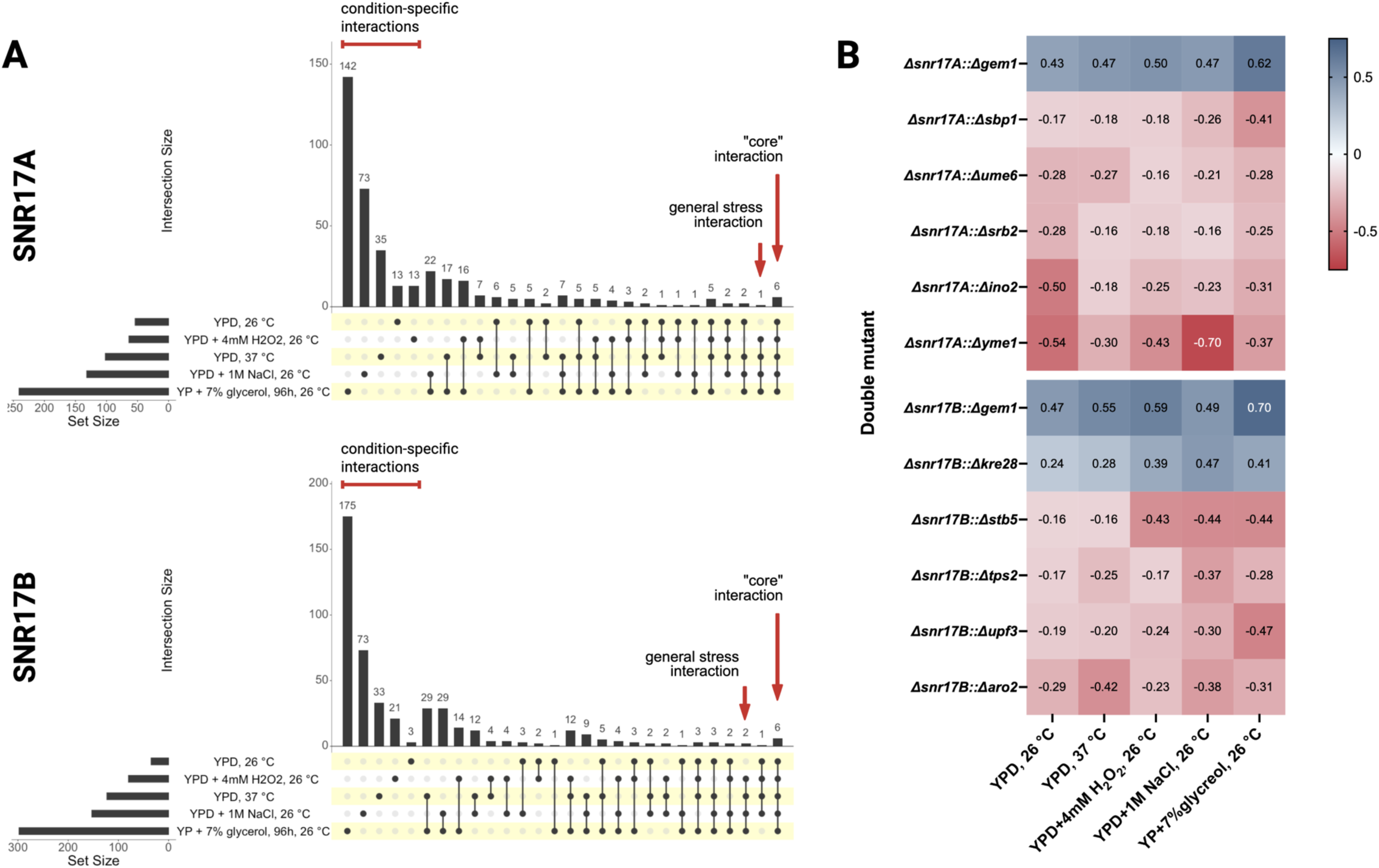
Environmental conditions elicit SNR17A and SNR17B interaction network changes. (**A**) Upset plot depicting environmental unique and shared genetic interactions between the five conditions tested (YPD, 26 °C; YPD + 4 mM H_2_O_2_, 26 °C; YPD, 37 °C; YPD + 1 M NaCl, 26 °C and YP + 7% glycerol, 26 °C) of SNR17A or SNR17B with coding genes. The bottom left-hand side horizontal bar chart displays the total number of significant interactions for each condition tested. The top vertical bar chart displays the number of interactions unique or shared between conditions indicated by a black dot in the bottom chart box. Unique interactions are represented by a single black dot between all 5 conditions tested. Several dots connected with a black line represent the number of interactions shared in the selected conditions. (**B**) Heatmap depicting genetic interactions that are shared among tested conditions. Six genetic interactions involving SNR17A and eight interactions involving SNR17B were shared across all five conditions tested. Specific interactions are arranged on the y-axis. Conditions are displayed on the x-axis. Negative interactions are represented as shades of red. Positive interactions are represented as shades of blue.

**Figure 9.**
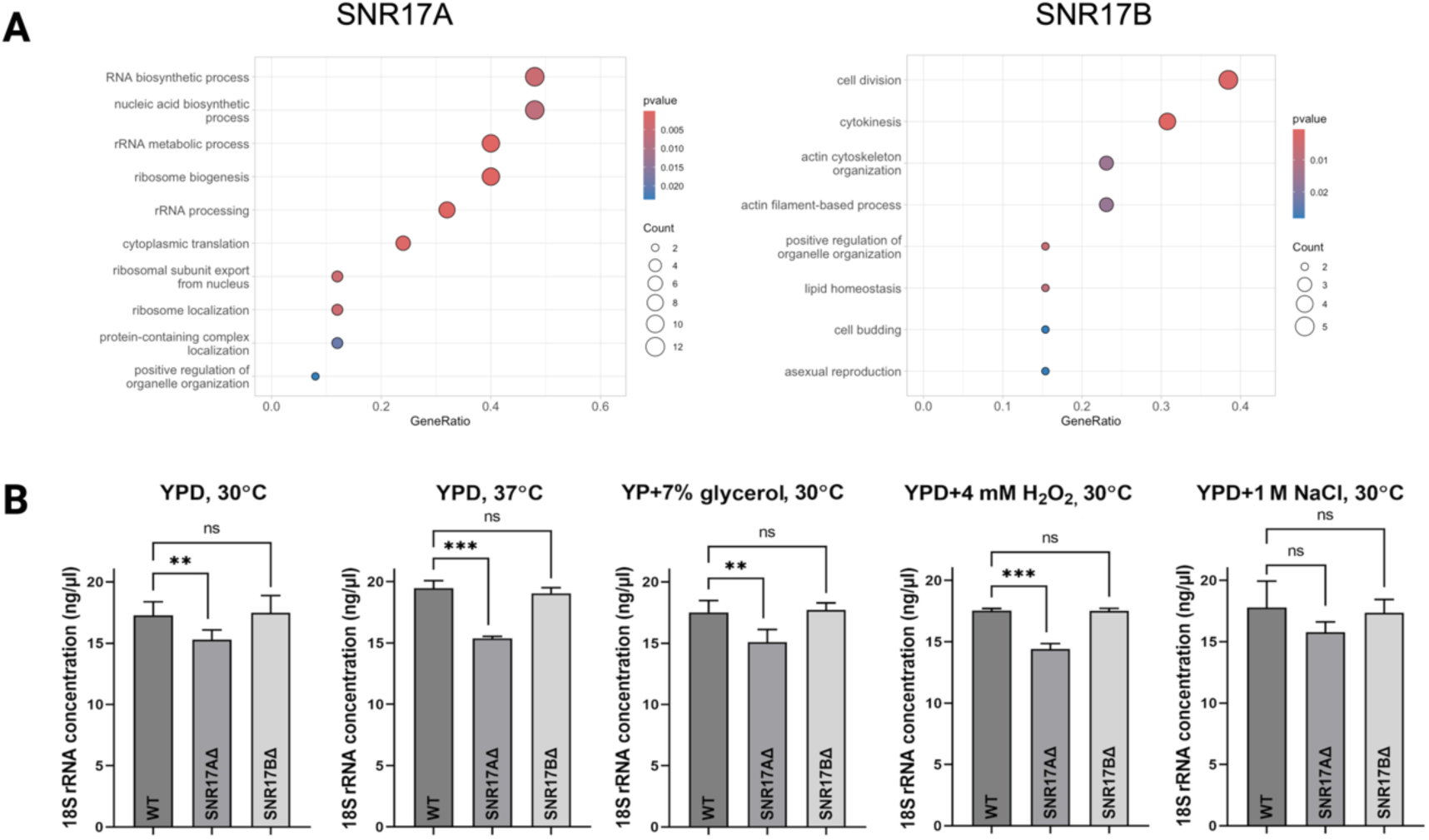
GO enrichment of SNR17A and SNR17B genetic interactions and 18S rRNA quantification. (**A**) GO enrichment of PGI with SNR17A or SNR17B in rich media. (**B**) Comparison of 18S concentration for the WT, SNR17AΔ and SNR17BΔ in 5 different conditions. Concentration of 18S rRNA in the WT and deletion mutants was analysed using the TapeStation 2200 system.

There were six and eight genetic interactions involving SNR17A and SNR17B, respectively, shared across all five conditions tested (Supplementary Dataset S10). These interactions maintain the same direction in all five conditions (Figure 8, panel B). This result echoes the “monochromatic” nature observed in the protein-coding network, where genetic interactions between defined pathways tend to be either exclusively positive or exclusively negative (Costanzo et al., 2010; Segrè et al., 2005). Among these interactions, three, *GEM1, SBP1,* and *UME6*, were common to both SNR17A and SNR17B, representing the core set of SNR17 snoRNA interactions conserved under all conditions. Beside *GEM1* that consistently exhibited strong positive interactions with both SNR17A and SNR17B as seen in rich media (Supplementary Dataset S9), *SBP1* (encoding nucleolar site-specific rRNA methyltransferase) and *UME6* (encoding Rpd3L histone deacetylase complex subunit) displayed negative interactions in all conditions (Park et al., 1992; Segal et al., 2006). Given that both *SBP1* and *UME6* are involved in translational (Segal et al., 2006) or transcriptional repression (Park et al., 1992), the synthetic sickness observed between U3 snoRNA and *SBP1* or *UME6* likely reflects an imbalance between ribosome biogenesis and gene expression, a relationship that appears to be conserved across all tested conditions.

Similarly, as was observed previously in the SNR17A and SNR17B ncRNAs networks (Figure 5), there is an expansion of the interaction network exclusive to SNR17B under the stress conditions, (Figure 6, panel B–E). Such a difference in the interaction network between SNR17A and SNR17B was consistent with what was observed in the ncRNA-ncRNA SGA, supporting the hypothesis of paralogs neo-functionalised.

We investigated the correlation between ΔSNR17 single-mutant query fitness and the number of their SGA interactions in each condition. No significant correlation was observed between query fitness and genetic interaction degree for either SNR17A (R² = 0.06) or SNR17B (R² = 0.007) (Supplementary Figure S6). This observation is consistent with the lack of correlation between the fitness changes in single mutants and the number of genetic interactions, as observed in the ncRNA-ncRNA SGA, supporting the idea that ncRNA interactions reflect broader functional dynamics within the cellular network.

### GO enrichment suggests a role for SNR17B in chromatin organisation under stress

Given the expansion of the SNR17B-exclusive interaction network under stress, which may indicate SNR17B stress specific neo-functionalization, we performed GO analysis on biological processes with the SGA data to assess the enrichment of GO terms among genes interacting with SNR17A or SNR17B (|ε| ≥ 0.15, p ≤ 0.001) to obtain their genetic interaction profiles and investigate insights into the respective functional roles of SNR17A and SNR17B.

We first focused on the GO analysis of positive genetic interactions (PGI) with SNR17A or SNR17B to identify physical interactions related to functional roles. Because U3 snoRNAs participate in rRNA processing by forming snoRNP complexes with proteins (*i.e*. through RNA-protein physical interactions), positive genetic interactions between SNR17 and rRNA-related proteins should be revealed by SGA as a reflection of physical interactions (Costanzo et al., 2016). For example, that is the case for *SLX9* interaction identified previously. Consequently, GO analysis of the PGI involving SNR17 paralogs should reveal enrichment in rRNA-associated pathways. As expected, rRNA-associated biological processes were enriched in SNR17A GO profiles in YPD 30 °C (**Error! Reference source not found.**, panel A) and in the stress conditions tested, except for YP + 7% glycerol, where no enrichment was observed (Supplementary Dataset S11). In contrast, no rRNA-related biological processes were enriched in the SNR17B PGI in YPD 30 °C (**Error! Reference source not found.**, panel A) or in any other conditions. These observations raised the hypothesis that SNR17B may have a different primary role in the cell than the rRNA-associated pathway and acts only secondarily as a backup of SNR17A. In fact, if SNR17B was truly redundant with SNR17A, it would be unlikely to have been maintained during evolution.

We measured the concentration of 18S rRNA in the WT and both deletion mutants grown in YPD and four stress conditions to investigate the involvement of SNR17A and SNR17B in rRNA processing. As SNR17 is crucial for 35S pre-rRNA processing and maturation of 18S rRNA (Maxwell & Fournier, 1995; Woolford & Baserga, 2013), the deletion of the U3 snoRNA paralogs is expected to reduce the concentration of the 18S rRNA fraction in the cell. As expected, the deletion of SNR17A resulted in a significant reduction of 18S rRNA abundance when compared to the WT strain in four out of five conditions tested (**Error! Reference source not found.**, panel B). This reduction in 18S rRNA concentration supports the role of SNR17A in pre-rRNA processing. In contrast, no significant changes in 18S rRNA abundance were observed with the SNR17B deletion mutant, suggesting that SNR17B is not the key player in maturation of 18S rRNA. Taken together, despite the high sequence similarity between SNR17A and SNR17B, the canonical U3-associated rRNA-processing function is carried out predominantly by SNR17A.

Given the hypothesis that SNR17B may have undergone neo-functionalization following the whole-genome duplication event, the question remains, what potential functions has SNR17B acquired? To answer this question, we analysed the GO profiles of SNR17A and SNR17B paralog-unique interactions under standard rich media, and compared these interactions with those acquired in the stress conditions. In rich media, GO analysis identified five significant GO terms unique to SNR17A interactions, one unique to SNR17B, and five shared between the two paralogs (Figure 10, panel A), which echoed the overall distribution of SNR17A and SNR17B genetic networks. In the stress conditions, SNR17A and SNR17B share only a limited number of GO terms. Specifically, a small overlap between the GO profiles involved stress response-related terms (Figure 10, panel B, C and D). In addition, the GO terms enriched for SNR17A and SNR17B under osmotic stress were entirely distinct (Figure 10, panel D). There is no enrichment in GO terms for both SNR17A and SNR17B in YP + 7% glycerol, indicating their genetic interaction in the obligatory respiratory condition is functionally dispersed across pathways and lacks the clustering for GO analysis. In addition, no enrichment of GO terms was observed for SNR17B under high temperature, showing that the expansion of SNR17B interaction in this condition could be a result of general broader stress response and not related to special functions. Taken together, the contrasting GO profiles enriched in the SNR17A- and SNR17B-specific interaction networks, particularly under stress conditions, provide direct functional evidence for divergence between the two U3 snoRNA paralogs.

**Figure 10.**
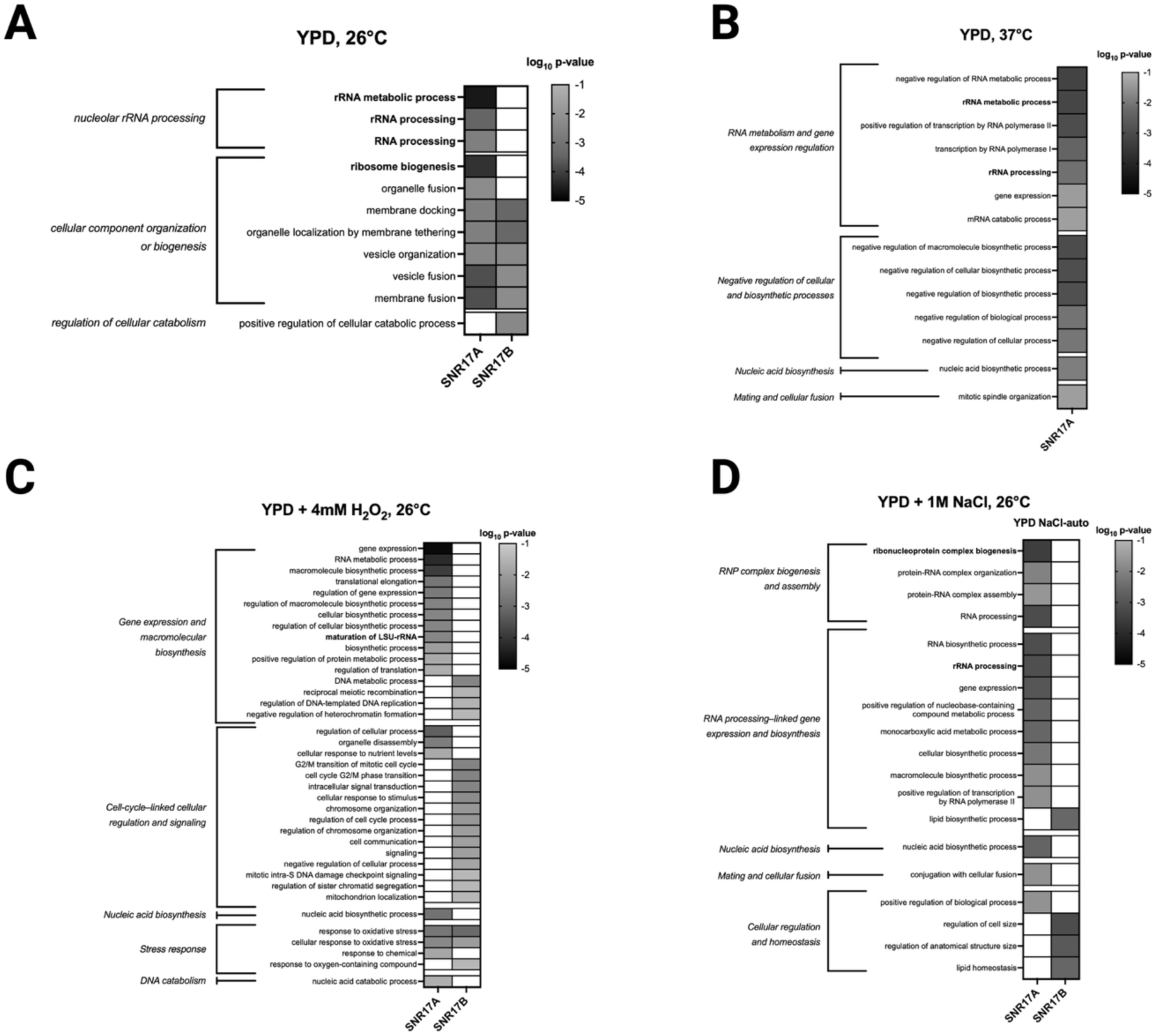
GO enrichment of interaction unique to SNR17A or SNR17B in each condition. **(A)**YPD, 26 °C. **(B)** YPD, 37 °C. **(C)** YPD + 4 mM H_2_O_2_, 26 °C. **(D)** YPD + 1 M NaCl, 26 °C. rRNA-associated biological processes are highlighted in bold.

SNR17B has GO-terms enriched in two of the four stress conditions tested, *i.e.* under oxidative stress and osmotic stress. In both conditions, we found that SNR17B exhibited significantly distinct and unique GO term clusters. When NaCl was present, the SNR17B GO profile was dominated by GO terms related to the regulation of cell size as well as lipid processing and homeostasis, whereas exposure to H₂O₂ led to enrichment of cell-cycle-associated categories (Figure 10, panel C and D). It has been reported that oxidative stress and high-salt conditions activate partially overlapping stress-response programs in yeast, including Hog1/Msn2–Msn4 signalling and mitochondrial redox regulation (Święciło, 2016). In addition, osmo-stimulated genes include proteins involved in protection against oxidative damage (Shirvanyan & Trchounian, 2024). Thus, SNR17B may serve a new function in response to the oxidative damage commonly caused by oxidative and osmotic stress.

To further investigate the neo-functionalization of SNR17B, we focused on the interactions that are unique to both oxidative and osmotic stress (*i.e.* interactions that are not in the other three conditions), which reflect the new function SNR17B gained specific to these two stresses. There are four interactions in common and unique for these two conditions including the genes *ICP55, RAD54, RPL11B* and *YSC83*. Interestingly, both *RAD54* and *RPL11B* have previously been linked to DNA conformation changes. *RAD54* encodes a motor protein of the SWI/SNF complex involved in homologous recombination and chromatin remodelling (Petukhova et al., 1999), whereas *RPL11B* has been reported to exhibit genetic interactions with chromatin-remodelling factors (Yadav et al., 2016). These interactors indicate that SNR17B associates with genes involved in chromatin structural remodelling and DNA topology. No function is known for *YSC83*, while *ICP55* is involved in mitochondrial protein stabilisation (Carrie et al., 2015)

Recently, some emerging evidence has implicated the U3 snoRNA in chromatin regulation beyond its canonical role in rRNA processing (Bratkovič et al., 2020). For example, in *Drosophila melanogaster*, a U3 snoRNA paralog has been shown to be required for the regulation of chromatin dynamics during viral infection (Jain et al., 2025). Notably, this snoRNA shares high sequence similarity with other U3 paralogs but contains a short, but distinct, RNA region absent from the other paralogs. This unique sequence enables the recruitment of chromatin remodellers to immune-related genes, thereby facilitating the stress-induced gene activation. Such paralog-specific acquisition of chromatin-associated regulatory function under stress echoes the divergence observed between SNR17A and SNR17B in yeast. In *S. cerevisiae*, DNA conformation changes are well established as a key regulatory mechanism underlying transcriptional responses to environmental stress, particularly oxidative and osmotic stress (Causton et al., 2001; Nadal-Ribelles et al., 2012). Under osmotic stress, for instance, the RSC complex, Hog1p, and Ssu72p are recruited to promote stress-induced gene looping at the CDC28 locus (Mendenhall & Hodge, 1998). Subsequent studies further demonstrated that the CDC28 antisense lncRNA plays an important role in this gene looping process by facilitating the docking of Hog1p to its target regions, thereby ensuring appropriate stress-responsive transcriptional regulation (Nadal-Ribelles et al., 2012, 2014). Given these established mechanisms, SNR17B may function in a similar mechanism, contributing to transcriptional regulation under oxidative and osmotic stress through modulation of chromatin dynamics.

## CONCLUSIONS

This study constitutes a pioneering effort to determine the genetic interactions involving 5 classes of ncRNAs, namely CUTs, SUTs, snoRNAs, tRNAs and RUF, and provides insights into their possible functions in yeast. In rich conditions, ncRNAs exhibit a higher proportion of positive genetic interactions, in contrast to the predominantly negative interactions observed within the protein-coding gene network (Costanzo et al., 2010). The networks of ncRNAs and their neighbouring genes are primarily independent, showing little overlap with each other. The small subsets of genetic networks that do overlap are composed predominantly of negative interactions, suggesting a potential effect *in cis* of the ncRNA on the flanking genes.

The environment has a significant impact on the ncRNA genetic interaction network. We observed an increase in the number of negative interactions in all stress conditions tested, with the highest number recorded in glycerol (up to 5.5 times more compared to rich media). This result suggests that these ncRNAs have unique functions in the cell or are being differentially transcribed depending on the nutritional context. The high plasticity of the ncRNAs genetic network suggests an important role in environmental adaptation.

The interaction data acquired in this study are also useful for distinguishing the cellular roles of ncRNA duplicates and for determining whether their functions have diverged over the course of evolution, as in the case of snoRNAs SNR17A and SNR17B. An analysis of SNR17A and SNR17B genetic networks in rich medium, revealed that these two paralogs exhibit most of the gene interactions, as expected. However, under stress conditions, SNR17B displayed a large number of unique epistatic interactions, suggesting that these paralogs may have neo-functionalised as a result of functional divergence that occurs after genome duplication and has acquired other roles in the cell.

Further analysis of the genetic interaction networks between SNR17 and protein-coding genes echoed the ncRNA-ncRNA results, revealing that SNR17B displayed a substantial number of paralog unique interactions under stress conditions. Through GO-term enrichment analysis and quantification of 18S rRNA levels in the SNR17A and SNR17B mutants, we found that SNR17A is a major player in rRNA processing, while SNR17B does not solely participate in this pathway despite its high sequence similarity to SNR17A, but has obtained a new function after the whole genome duplication event. Additional GO analysis of the stress-specific SNR17B interactions further suggested a potential neo-functionalization associated with DNA conformation change that potentially facilitates transcriptional regulation under oxidative and osmotic stress.

Overall, this study offers the first insight into the environmentally-dependent epistatic interactions of ncRNAs in a eukaryotic organism.

## MATERIALS AND METHODS

### Strains and plasmids

We previously generated a library of ncRNA mutants in both haploid and diploid backgrounds (Parker et al., 2017, 2018). The *MAT***a** library from this collection (Supplementary Dataset S2), referred here as ‘array’, was used in the crossing with the haploid *MATα* ‘query’ single deletion strains (generated in this study; see Supplementary Materials and Methods and Supplementary Dataset S1) to create double mutants according to the SGA protocol (Baryshnikova et al., 2010). The *MAT***a** array library was constructed in the BY4741 background (*his3Δ1 leu2Δ0 met15Δ0 ura3Δ0*) and all these mutants carry a *kanMX* resistance gene that replaced the ncRNAs of interest (Parker et al., 2017). The *MATα* query collection was generated in the Y7092 background (*can1Δ::STE2pr-Sp_his5 lyp1Δ ura3Δ0 leu2Δ0 his3Δ1 met15Δ0*; kindly provided by the Boone lab) (Tong & Boone, 2007). The query ncRNA deletion strains carry the *natMX4* resistance marker that replaced the ncRNAs of interest. This *natMX4* marker confers clonNAT (Nourseothricin; WERNER BioAgents GmbH) antibiotic resistance and was amplified from the pFA6*-natMX4* (Wach et al., 1994) plasmid in a single PCR step using primers partly complementary to the ncRNA flanking sequences.

### Query strain library construction, SGA and quality control of double mutants generated

The query strains were constructed by substituting the ncRNA loci with the *natMX4* cassettes. Each cassette was amplified using a pair of barcoded primers containing genome-complementary sequences from pFA6-*natMX4* (Chu & Davis, 2008) (see Supplementary Figure S7 and Supplementary Materials and Methods). The list of all query strains constructed in this study can be found in Supplementary Dataset S1. Thirty-four query strains were used to generate a double ncRNA mutant library (see Supplementary Materials and Methods). The SGA was performed three times using three biological replicas of the query strains. Following SGA, images of the final selection plates containing generated double ncRNA mutants were recorded in white light using a Phenobooth and cropped using Corel PaintShop Pro X7. Subsequently, the images were processed using the high-throughput SGA data analysis software *Balony* (Young & Loewen, 2013) that measures colony area in pixels.

Each SGA array has technical replicas of both array mutants (Supplementary Dataset S2) and double mutants (Supplementary Dataset S3). Some ncRNAs mutants have four and others eight replicas, so for each genetic interaction we have at least 12 data points that allowed for a robust analysis.

Furthermore, following SGA, 20 randomly selected haploid ncRNA double mutants were validated by PCR (see Supplementary Materials and Methods, Supplementary Figure S8, Supplementary Table S1).

In addition to double ncRNA mutants, ncRNA-protein coding double mutants between SNR17A&B and protein deletion collection were generated using SGA (Baryshnikova et al., 2010). A subset of non-essential yeast protein deletion collections, namely the diagnostic collection (Kuzmin et al., 2021) was used as the array. The array was crossed with either SNR17A or SNR17B query strains to generate a library of ncRNA-protein double deletion mutants. All experiments were performed at the permissive temperature of 26 °C to maintain the viability of the temperature-sensitive mutants.

After generating ncRNA-protein double mutants, an additional quality control step was applied to verify synthetic lethal interactions before conducting phenotypic analyses. This extra validation step aimed to minimise false positives arising from technical issues, such as the failure to pick up low-fitness colonies by RoToR pinpads. All double mutants underwent reselection from sporulation plates, followed by an extended incubation period to allow sufficient cell mass for plate replication. Only double mutants that exhibited growth on all plates except the final selection plate were considered true synthetic lethal candidates and retained for further analysis. Following SGA, 5 randomly selected double mutants were validated by PCR.

### SGA data analysis and identification of epistatic interaction

The initial normalisation was performed in this software to correct for uneven plate growth according to the plate median and the row/column correction was applied. Minimum and maximum spot sizes were set to 0.02 and 100, respectively. Following the initial image processing in the Balony software, the generated data was analysed using the R package version 3.4.4 (R Core Team, 2023). The initial analysis assessed the growth consistency of the double mutants created through the SGA. The data was filtered to remove ***i.*** single and double ncRNA mutants that showed inconsistent growth patterns among the quadruplicates/octuplicates and ***ii.*** linkage group of ncRNAs that were on the same chromosome and were scored as lethal after the SGA as their phenotype was likely to be linked to the lack of meiotic crossing-over between them.

The data was processed using an in-house R script. Recorded output size values for single and double mutant colonies were used to generate an SGA score epsilon as described in (Baryshnikova et al., 2010) and (Wagih et al., 2013).

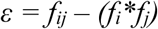

where f_i_ and f_j_ are the fitnesses of the single mutants carrying mutations of genes *i* and *j* respectively, and f_ij_ the double mutant fitness. An estimated query fitness was applied to calculate epsilon as described in (Baryshnikova et al., 2010). It is assumed here that the relative size defect of the double mutant colony (f_ij_) compared to expectation (f_i_f_j_) relates to the effect of genetic linkage or genetic interaction. In the absence of genetic linkage, just like in the absence of genetic interactions, the fitness of a double mutant should equal the product of the two single mutants’ fitness.

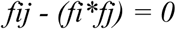

A score −0.15 ≥ *ε* ≥ 0.15 represents no change in growth, and thus no genetic interaction. A score ≥ 0.15 would represent an increase in fitness, and thus a positive genetic interaction. A score of −0.15 would represent a decrease in fitness in the double mutant compared to the single mutant expectation, and thus a negative genetic interaction. Double mutants in that case were described as having a synthetic sick phenotype, but if the double mutant fitness f_ij_ was ≤ 0.07, the phenotype was described as “lethal”.

For SGAs involving protein-coding gene mutants, an additional correction was applied to the double mutant fitness for each plate by adjusting the epsilon values of *HIS3::Kan* array strains with any query to zero, in order to avoid bias toward positive interactions caused by the overall low fitness of these double mutants.

We used Z-scores, which are standard deviations, to calculate the p-values of the samples. A sample here is an SGA score. First, we assumed a normal distribution. The p-value was calculated for each sample mean, μx ,with x being the SGA score. Here, the p-value is the probability that we would obtain a given mean a that is greater than the absolute value of its Z-score or less than the negative of the absolute value of its Z-score. The null hypothesis is that a given mean is equal to the absolute value of its Z-score.

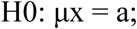

The Z-score was found by assuming that the null hypothesis is true, subtracting the assumed mean, and dividing by the theoretical standard deviation. Once the Z-score is found, the probability that the value could be less the Z-score is found using the pnorm R command. But this is not enough to get the p-value. If the Z-score that is found is positive then we need to take one minus the associated probability. Also, for a two-sided test we need to multiply the result by two. Here, we avoid these issues and ensure that the Z-score is negative by taking the negative of the absolute value. That is why our formula in our script is:

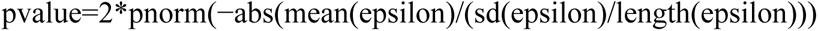

Assuming a normal distribution, p-values were calculated for each sample mean using the Z-score. The Z-score was found by assuming that the null hypothesis was true, subtracting the assumed mean, and dividing by the theoretical standard deviation. Q-values, which are Bonferroni-adjusted p-values, were inferred using the “stats” R package (R Core Team, 2023). We selected as significant epistasis the cases where the double mutants had an SGA score |ε| ≥ 0.15 (as previously adopted by Costanzo et al., 2016) associated with a stringent q-value ≤ 0.001. The “UpSetR” R and “matplotlib_venn” Python packages were used for data visualisation.

### SGA network generation

Genomic interaction networks were created from the data generated for all experimental conditions tested. These included yeasts grown in rich media as well as on non-fermentable media, oxidative, high temperature and osmotic stress environments. The networks were generated using the “visNetwork” and “igraph” R libraries (Almende et al., 2025; Csárdi & Nepusz, 2006). Nodes represent the ncRNAs, and the edges represent the genetic interaction between them. The size of the nodes is proportional to the number of connections with the other nodes. The edges are coloured in green and red to distinguish the positive and negative epistasis, respectively. The thickness of the edges is proportional to the absolute value of the fold change. The force-directed layout (forceAtlas2Based solver) (Jacomy et al., 2014) has been used to organise the graph in a 2D space. The central gravity of the networks is distance independent. The repulsion between the nodes is linear.

### Phenotypic analysis of the mutants in stress conditions

The phenotypic analysis of both double and single deletion strains was performed in five different environmental conditions on solid growth media.

For ncRNA-ncRNA SGA, the condition included (i) yeast rich media, YPD agar at 30 °C, and four stress conditions: (ii) high temperature on YPD agar at 37 °C, (iii) obligatory respiratory on YP agar supplemented with 7% glycerol at 30 °C, (iv) osmotic stress on YPD agar supplemented with 1 M NaCl at 30 °C and (v) oxidative stress on YPD agar supplemented with 4 mM H_2_O_2_ at 30 °C.

The single and double deletion library was analysed in the same 1,536 format as described for SGA above. All cells were first grown on solid YPD agar media for 48 h prior to the experiment and subsequently transferred to the specific experimental plates using the RoToR HDA system and grown for 48 h at their relevant temperatures before analyses. The images of these plates were recorded using a Phenobooth and analysed using the Balony software in a similar manner as described for the SGA above. We used the same cut-off analysis values as described above for the SGA analysis in standard yeast condition (|ε| ≥ 0.15 and q-value ≤ 0.001).

For ncRNA-protein SGA, phenotypic analysis of both double and single deletion strains was performed in the previously mentioned five environmental conditions, with a minor modification of the growth temperature from 30 °C to 26 °C, except for the high-temperature condition, which remained at 37 °C on YPD agar. Plate images were captured using a Phenobooth (Singer Instruments, UK) after 96 h for YP + 7% glycerol and after 48 h for the remaining four conditions.

### Genetic and physical interactions between neighbouring genes/proteins of interacting ncRNAs

Protein-protein physical interactions as well as genetic interactions between coding genes that were located in the neighbourhood of every interacting ncRNA pairs were analysed. Only significantly interacting ncRNA pairs were included. A synthetic genome annotation file (BED file format) containing the names of protein-coding genes and ncRNAs, their positions and strand information, was generated using an in-house Python 3.6 script. Gene annotations were extracted from the *Saccharomyces* genome database (SGD; www.yeastgenome.org/) and ncRNA annotations were added manually based on the information from Wery et al., 2016. Subsequently, the genes and ncRNA were ordered by their start position and by the chromosome number using the BedTools sort tool (Quinlan & Hall, 2010). The flanking genes for each ncRNA were extracted using an in-house Python 3.6 script, generating 4 combinations of gene interactions for each pair of interacting double ncRNAs. A Python 3.6 script that uses the Representational State Transfer (REST) Application Programming Interface (API) of BioGRID database (https://thebiogrid.org/) was developed to query and access its data on the protein coding gene interaction data (physical and genetic) as well as phenotype observed and experiment type (Oughtred et al., 2019; Stark et al., 2006). Multiple hits of the same type of interaction (*i.e.* physical or genetic) with different approaches were counted only once for each coding gene combination. The null expectation for how often the protein-coding neighbours should interact, *i.e*. P(H_0_), was calculated by total recorded number of PPI vs. total combination of gene pairs.

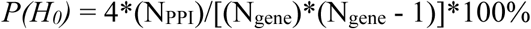

Where N_PPI_ represents the total number of recorded non-redundant PPIs in BioGRID, and N_gene_ is the total number of genes in *S. cerevisiae* strain S288C.

Pearson’s correlation was used to measure associations between the number of interactions and single mutant fitness defect, and the Pearson correlation coefficient (r) was calculated.

### Growth analysis of SNR17AΔ and SNR17BΔ single and double mutants

A spot growth assay was performed on the single SNR17AΔ and SNR17BΔ mutants in the yeast standard media to reveal phenotypic effects of the snoRNA deletion. At least two biological replicates were tested for each mutant. The growth was compared to the host WT strain Y7092. Cells for the spot assay were prepared in the following way: cells were grown overnight with shaking (200 rpm) in YPD at 30 °C, and the cultures were centrifuged for 2 min at 2000 rpm. The pellet was resuspended in YPD and diluted to a starting OD_600_ of 1.0. Two-times or ten-times dilutions were prepared and 1 and 5 µL of the cell suspensions were spotted on an agar plate, which was incubated for 48 h at 30 °C or 37 °C, depending on the environmental condition. The growth was recorded every 24 h of incubation.

### Gene Ontology enrichment analysis

Enrichment analysis was performed for SNR17A and SNR17B SGA genetic interaction using the R package clusterProfiler (Yu et al., 2012) which included Gene Ontology (GO) terms. GO terms with p ≤ 0.05 were considered significantly enriched. The analysis encompassed the Biological Process (BP) ontology. A total of 1,032 array mutant gene strains served as the background for calculating enrichment p-values.

GO results were simplified using the Wang semantic similarity method and visualised with ggplot2. The simplifyEnrichment package (Gu & Hübschmann, 2023) was used to cluster enriched GO terms based on the binary cut algorithm. Heatmaps of clustered GO terms were generated using GraphPad Prism (ver. 10.6.0).

## ACKNOWLEDGMENTS

This work was supported by the Wellcome Trust, under the grant number 094225 to RTO and DD and 104981 to RTO, SGJ and DD. ST was supported by the European Research Council, H2020-MSCA-ITN-2017, under the grant number 764364 awarded to DD (https://cordis.europa.eu/project/id/764364) and by the Future Biomanufacturing Research Hub (Future BRH), funded by the Engineering and Physical Sciences Research Council (EPSRC) and Biotechnology and Biological Sciences Research Council (BBSRC) as part of UK Research and Innovation (grant EP/S01778X/1). LNB was supported by The National Secretary of Higher Education, Science, Technology and Innovation (SENESCYT, http://siau.senescyt.gob.ec/), Ecuador, and by the BBSRC grant BB/T002123/1 awarded to DD. KD scholarship was funded by the Royal Government of Thailand under the scheme for Development and Promotion of Science and Technology Talent (DPST) projects. MWF was supported by the University of Manchester - China Scholarship Council joint scholarship. The authors wish to thank Charlie Boone, Michael Costanzo and Sondra Bahr for useful discussions, provision of the Y7092 strain and help with the SGA methodology. The authors also thank Diego Estrada-Rivadeneyra for initial help with U3 snoRNA data acquisition.

## DATA AVAILABILITY

The interactive ncRNAs networks have been added to the publicly accessible online resource at https://github.com/DelneriLab/SGA_ncRNA_project.

## CODE AVAILABILITY

Codes used to analyse the data in this manuscript are available at https://github.com/DelneriLab/SGA_ncRNA_project.

