## Supplementary Figures and Methods for "Genome-wide analysis of the non-coding RNA synthetic genetic network reveals extensive plasticity and distinct environmental dependent roles for U3 snoRNA paralogs"

**Supplementary information**

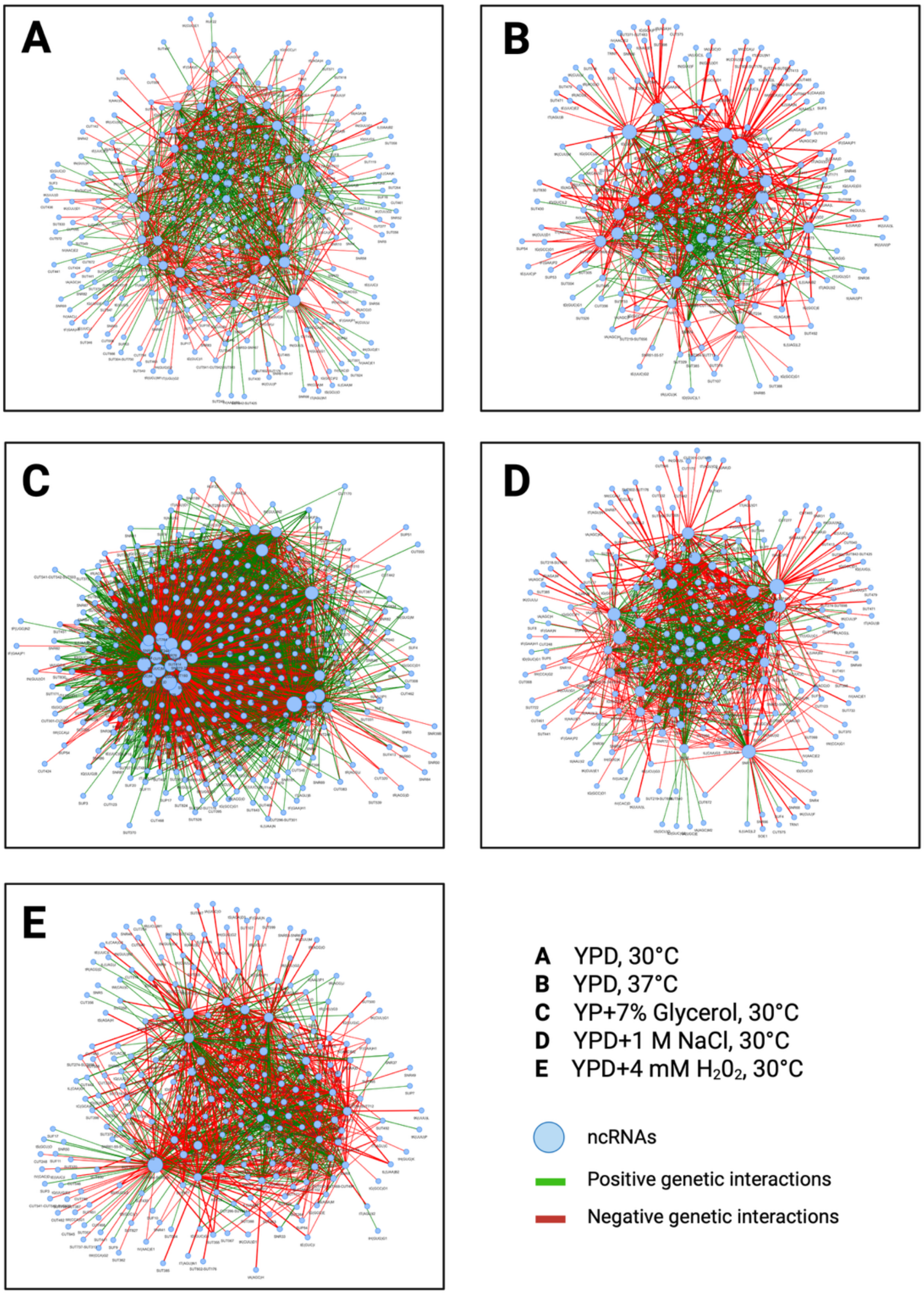


**Supplementary Figure S1. Visualisation of the ncRNA network under different environmental conditions.** ncRNA network generated for 29 query ncRNA deletion strains based on the $|\varepsilon| \geq 0.15$ and q-value $\leq 0.001$ cut off in (A) YPD, 30 °C, (B) YPD, 37 °C, (C) YP + 7% Glycerol, 30 °C, (D) YPD + 1 M NaCl, 30 °C and (E) YPD + 4 mM H2O2, 30 °C. Network ncRNAs nodes (blue); negative ncRNA genetic interactions (red edges); positive ncRNA genetic interactions (green edges). The thickness of the edges is proportional to the absolute epsilon.

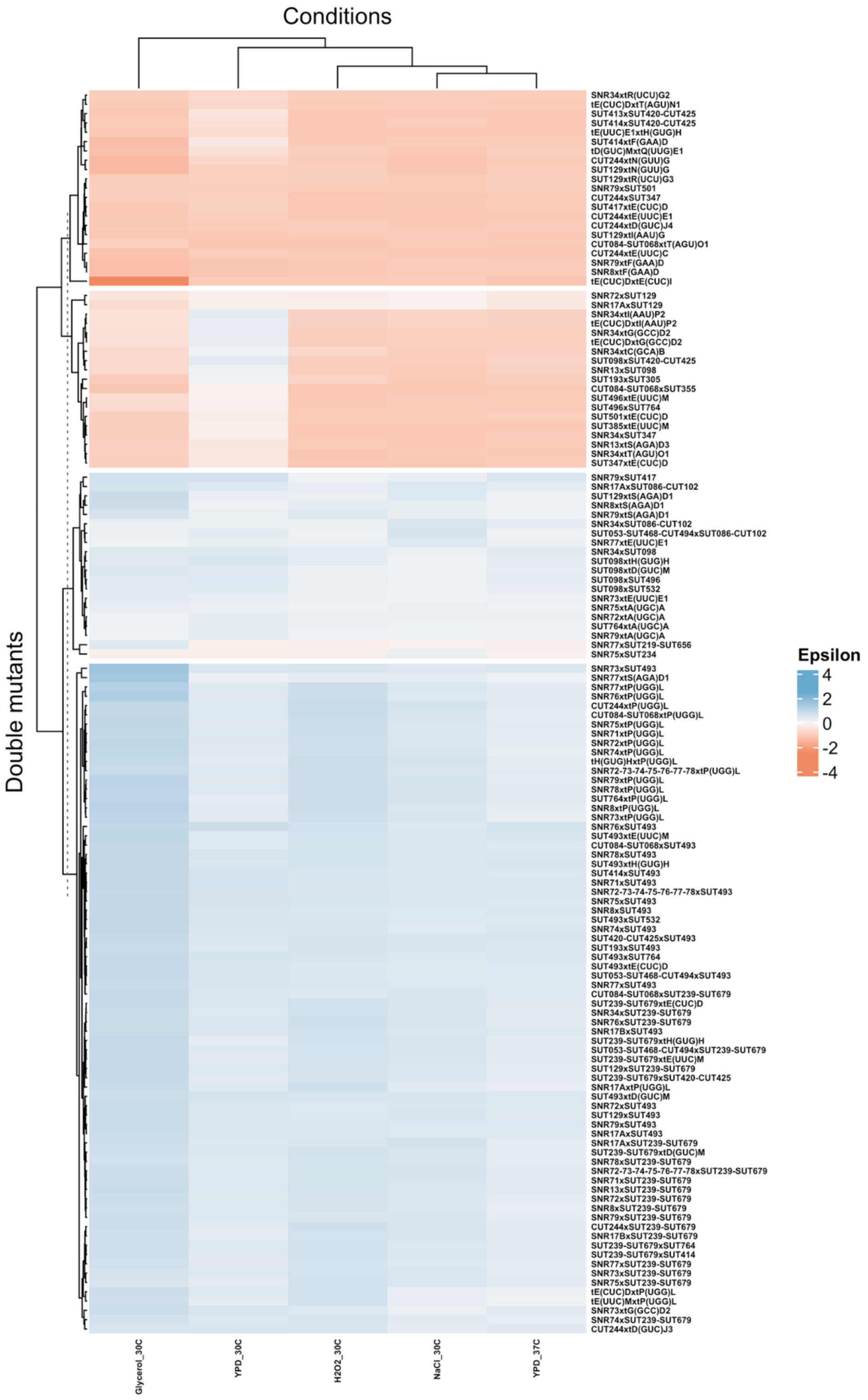


**Supplementary Figure S2.** Heatmap depicting epsilon of 130 “core ”ncRNA interaction pairs that were shared between all five conditions tested. ncRNA pairs are arranged on the y-axis, and clustered using hierarchical clustering. Conditions are displayed on the x-axis. Negative interactions are represented as red and positive interactions are represented as blue.

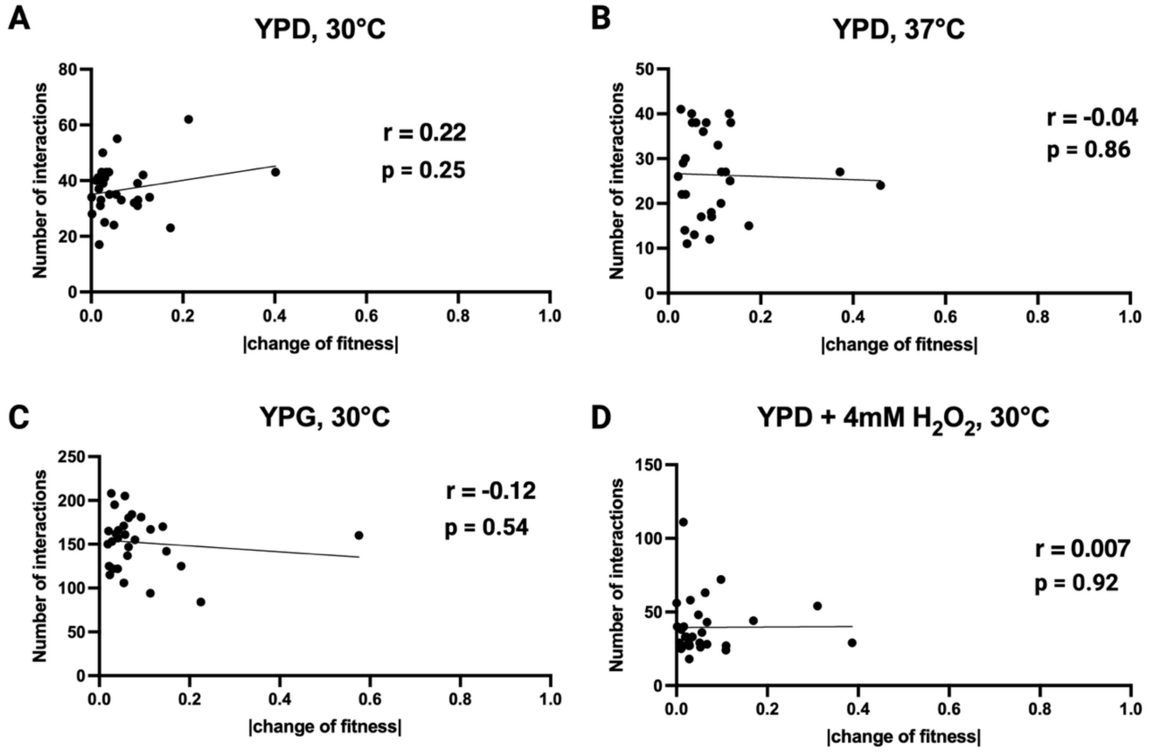


**Supplementary Figure S3.** Scatter plot displaying Pearson's correlation between the absolute fitness changes and the number of interactions. Number of interactions were correlated with the fitness changes in (**A**) YPD, 30 °C, (**B**) YPD, 37 °C, (**C**) YP + 7% glycerol, 30 °C and (**D**) YPD + 4 mM $H_2O_2$, 30 °C.

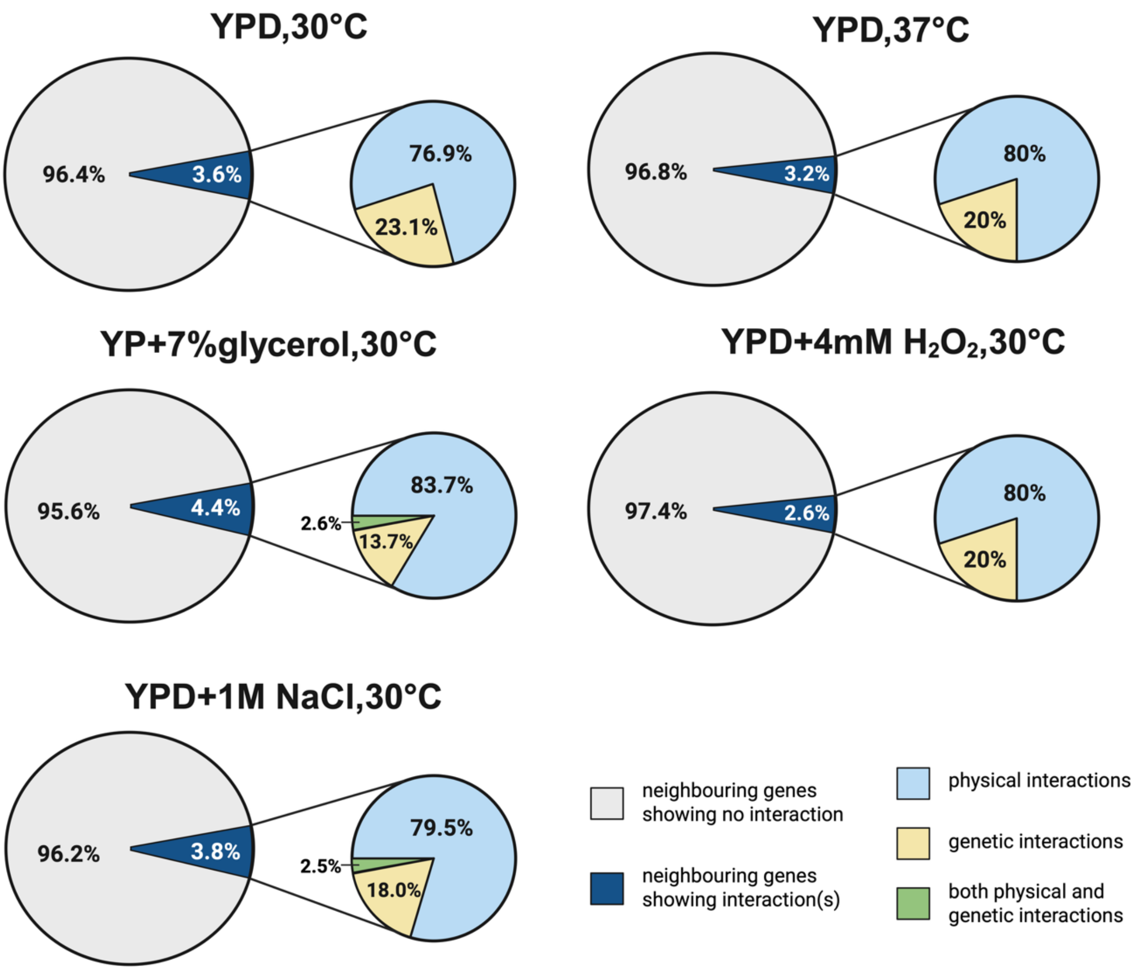


**Supplementary Figure S4.** Genetic and physical interactions in environmental conditions. Pie chart representation of genetic and physical interactions analysed between the neighbouring genes of the significantly interacting ncRNAs**.** Dark blue colour in the larger pie charts represents the percentage of ncRNAs protein-coding neighbouring genes with a significant type of interactions. Smaller pie charts display the proportion of these interactions broken down into genetic (yellow), physical interactions (light blue) or both interactions (green) in YPD, 30 °C; YPD, 37 °C; YP+ 7% glycerol, 30 °C; YPD + 1 M NaCl, 30 °C; and YPD + 4 mM $H_2O_2$, 30 °C.

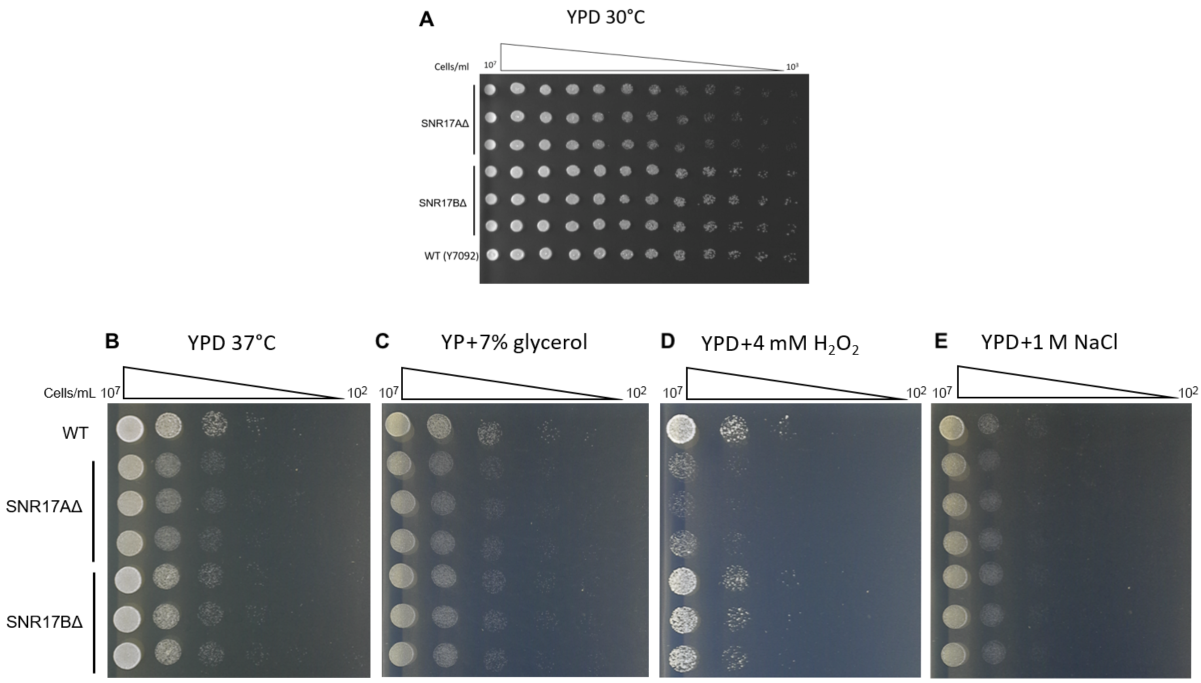


**Supplementary Figure S5.** Spot assay to determine growth differences between SNR17A and SNR17B deletion strains. Serial dilutions (2x for **A**, and 10x for **B** to **E**) of three biological replicates of haploid SNR17AΔ and SNR17BΔ mutants along with a host strain Y7092 were spotted on YPD, 30°C (**A**), YPD, 37°C (**B**), YP + 7% glycerol, 30°C (**C**), YPD + 4 mM $H_2O_2$, 30°C (**D**) and YPD+1 M NaCl, 30°C (**E**). Growth was recorded after 24h.

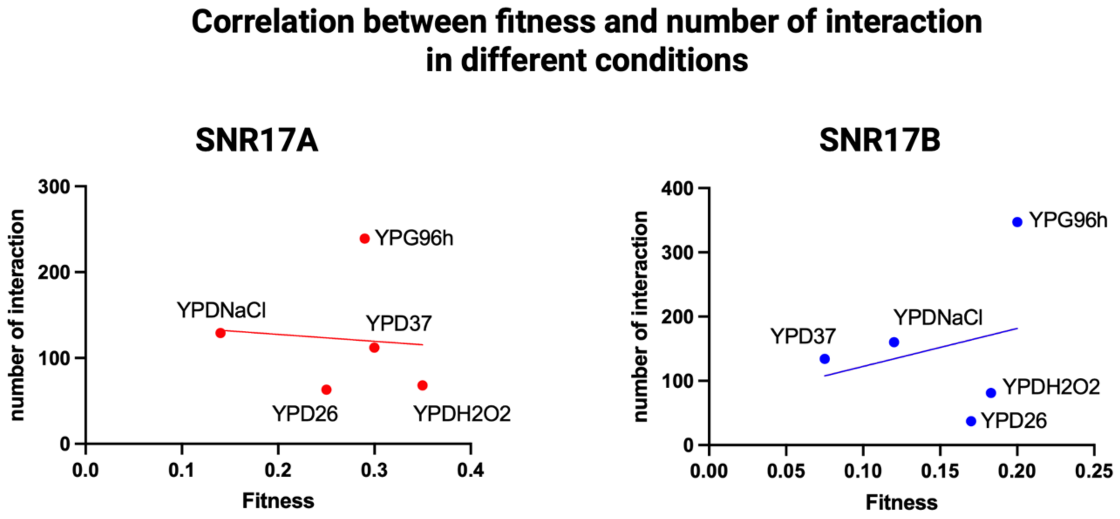


**Supplementary Figure S6.** Scatter plot showing the Pearson correlation between absolute fitness changes and the number of genetic interactions for SNR17A (red) and SNR17B (blue). Correlations were assessed across five tested conditions: YPD at 26 °C, YPD at 37 °C, YP + 7% glycerol at 26 °C, YPD + 4 mM $H_2O_2$ at 26 °C, and YPD + 1 M NaCl at 26 °C. Each point represents one condition. The x-axis indicates the absolute fitness change of the query ncRNA mutant (*i.e.* SNR17A or SNR17B) in the corresponding condition, and the y-axis indicates the number of interactions.

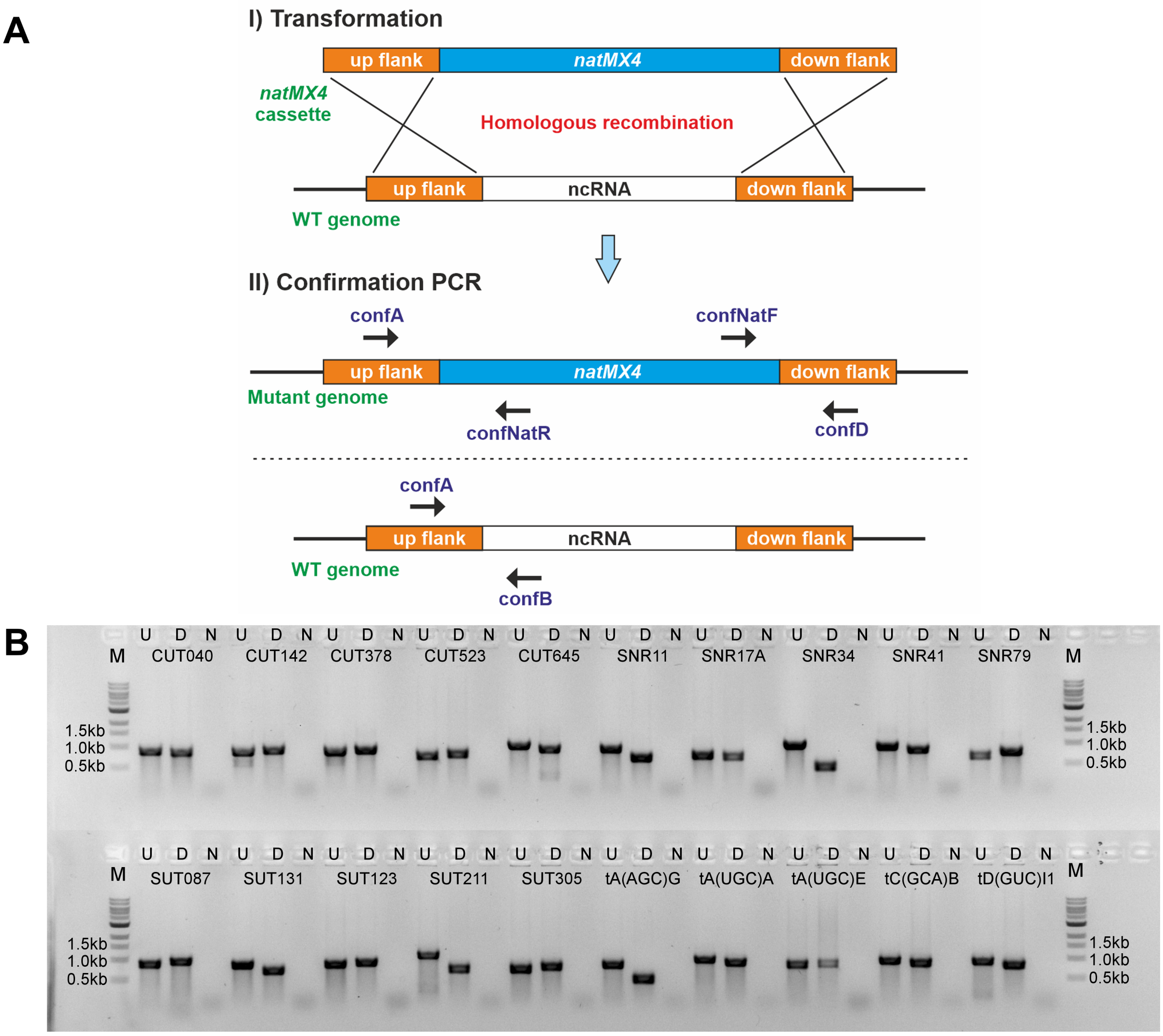


**Supplementary Figure S7.** Construction and validation of query strains. (**A**) Strategy to construct and validate ncRNA query deletion mutants. Briefly, ncRNA deletion cassette composed of the natMX4 marker and ncRNA complementary flanking regions were transformed into the MAT*α* strain (Y7092) to delete the ncRNAs of interest. Following transformation, single colonies were PCR validated with primer sets: (i) confA + confNatR, (ii) confNatF + confD and (iii) confA + confB. Primer sets (i) and (ii) only amplified PCR bands if the deletion cassettes were integrated at the correct loci. Primer set (iii) amplified a PCR band if no integration event occurred. (**B**) Agarose gel electrophoresis for the PCR validation of randomly selected ncRNA query mutants. U: upstream flank PCR with primer set confA + confNatR; D: downstream flank PCR with primer set confNatF + confD; N: control PCR for the native ncRNA, with primer set confA + confB; M: 1 kb New England Biolabs ladder.

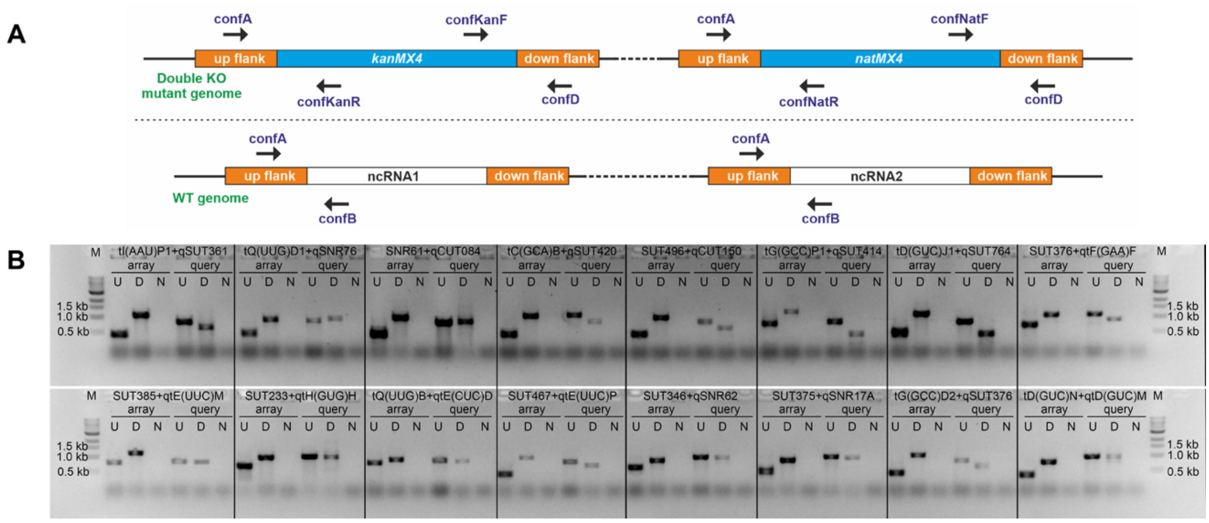


**Supplementary Figure S8.** Validation of a subset of SGA double deletion mutants. (**A**) Strategy to validate the double ncRNA deletion mutants. Following SGA, randomly selected double deletion haploid mutants were subjected to confirmation PCR. Primer sets (i) confA + confKanR and (ii) confKanF + confD were used to test for the *kanMX4* deletion cassette integrations. Primer sets (iii) confA + confNatR and (iv) confD + confNatF were used to test for the *natMX4* deletion cassette integrations. Primers confA + confB provided additional ncRNA deletion confirmation, only amplifying bands if the ncRNAs were intact. (**B**) Agarose gel electrophoresis for PCR validation on a selection of mutants. U: upstream flank PCR with primer set confA + kanB or confNatR; D: downstream flank PCR with primer set kanC or confNatF + confD; N: control PCR for the native ncRNA with primer set confA + confB. M: 1 kb New England Biolabs ladder.

**Supplementary Datasets**

- **Supplementary Dataset S1** - Constructed query ncRNA strains
- **Supplementary Dataset S2** - Array ncRNA mutants
- **Supplementary Dataset S3** - Double KO significant epistasis
- **Supplementary Dataset S4** - Genetic-physical interactions YPD
- **Supplementary Dataset S5** - Number of significant ncRNA interactions different condition and epsilon
- **Supplementary Dataset S6** - Individual and shared epistatic interactions.001
- **Supplementary Dataset S7** - Genetic-physical interactions, various conditions
- **Supplementary Dataset S8** - SNR17A&B interactions ncRNA SGA
- **Supplementary Dataset S9** - SNR17A&B interactions ncRNA SGA
- **Supplementary Dataset S10** - Shared and unique interaction in different conditions
- **Supplementary Dataset S11** - GO analysis of PGI

### Supplementary Materials and Methods

#### Query strain library construction

The query strains were constructed by substituting the ncRNA loci with the *natMX4* cassettes. Each cassette was amplified using a pair of barcoded primers containing genome complementary sequences (Chu & Davis, 2008) from pFA6-*natMX4* in the total of 25 µl PCR volume: 10 ng plasmid DNA, 5 µM of forward and 5 µM of reverse primers, 10 mM of each dNTP, 2.5 units of LongAmp polymerase (New England Biolabs) and 1x of reaction buffer. The cycling conditions were as follows: initial denaturation at 95°C for 2 min followed by 35 cycles of 94°C for 30 sec, 58°C for 30 sec and 65°C for 1.5 min and 1 step of final elongation of 65°C for 5 min. The amplified PCR products were ~1.5 kb and they were transformed into the Y7092 (*MATα*) background strain. Transformants were selected on YPD agar (1% yeast extract, 2% peptone, 2% dextrose, 2% agar) supplemented with 200 µg/ml of clonNAT (Werner BioAgents, Jena, Germany). After 4 days of growth at 30°C, 1 to 5 colonies were streaked, and single colonies were confirmed by colony PCR for correct cassette integration. The list of all query strains constructed in this study can be found in Supplementary Dataset S1. Thirty-four query strains were used to generate double ncRNA mutant library.

#### PCR confirmation of query strains

Correct ncRNA locus deletion was confirmed by colony PCR. Following transformation, streaked colonies were resuspended in 100 µl sterile $H_2O$. Three sets of primers were used: (i) confA-confNatR, (ii) confNatF-confD and (iii) confA-confB (Supplementary Figure S7). Confirmation primers confA, confB and confD are listed in Parker et al. (2017). Primer pairs (i) and (ii) only generated PCR products if the ncRNA was replaced with the deletion cassette. Primer set (iii) generated PCR products if the ncRNA was intact and these colonies were discarded. The PCR was performed in the total volume of 25 µl and contained 5 µl of resuspended cells, 5 µM of forward and 5 µM of reverse primers, 10 mM of each dNTP, 2.5 units of LongAmp and 1x of reaction buffer. The same PCR conditions as above were used, generating PCR products between 0.6 kb and 1.2 kb, depending on the deleted ncRNA, which were analysed on a 1.5% agarose gel. Confirmed colonies were stored in YPD containing 15% glycerol at -80C until needed.

#### Quality control of the double mutants created using SGA

Following SGA, 20 randomly selected haploid ncRNA double mutants were validated by PCR. The following primer sets were used for ncRNA1::*kanMX4* validation: (i) confA+confKanR,

(ii) confD+confKanF and (iii) confA+confB. Primer sets (i) confA+confNatR, (ii) confD+confNatF and (iii) confA+confB were used for ncRNA2::*natMX4* validation in the same mutant (Supplementary Figure S8).

**Single deletion ncRNA library formatting for SGA**

The single deletion array library (*MATa*; ncRNA*::kanMX4*) is composed of 376 mutants encompassing including 44 CUTs, 92 SUTs, 58 snoRNAs, 181 tRNAs and 1RUF22 (Supplementary Dataset S2) (Parker et al., 2017). The library was first organised in 96 and 384 density formats and subsequently used as a source to generate the final working copy of two Plus Plates (Singer Instruments, UK) in a density of 1536 colonies per plate using the RoToR HDA system (Singer Instruments, UK). Each single deletion strain was plated in quadruplicates allowing replicates for consistent SGA analysis. The edges of each plate were excluded from the SGA analyses since colonies that are located on the edges and corners have more nutrient availability and decreased competition with the neighbouring colonies and hence often show increased growth (Baryshnikova et al., 2010b). To minimise contamination, the single deletion ncRNA plates were grown on YPD agar supplemented with 200 µg/ml G418 (Sigma-Aldrich) or SD complete amino acid agar (0.67% yeast nitrogen base with amino acids (Merck), 2% glucose, 2% agar) supplemented with 200 µg/ml G418. The images of these plates were recorded using a PhenoBooth (Singer Instruments, UK) for subsequent SGA analyses.

**Generation of the double ncRNA mutants using SGA**

Thirty-four query ncRNA deletion mutant strains (*MATα*; ncRNA*::natMX4*) (Supplementary Dataset S1) were selected to generate a library of double ncRNA deletion mutants. Prior to SGA, a small portion of -80°C cell cultures of these query strains were grown overnight in 5 ml of liquid YPD at 30°C with shaking (200 rpm). Subsequently, 1 ml of the overnight cultures was spread onto PlusPlates containing YPD supplemented with 200 µg/ml of clonNAT using plastic cell spreaders and grown overnight at 30°C. Then, the query strains were crossed with the array ncRNA deletion library in the 1536 format on YPD agar using a RoToR HDA robot and disposable plastic replicators (pinning RePads, Singer Instruments, UK). SGA was carried out according to Baryshnikova et al. (2010a) including the generation of the double ncRNA mutant library and subsequent analysis of epistasis.

**References to Supplementary Materials and Methods**


Balarezo-Cisneros, L. N., Parker, S., Fraczek, M. G., Timouma, S., Wang, P., O'Keefe, R. T., Millar, C. B., & Delneri, D. (2021). Functional and transcriptional profiling of non-coding RNAs in yeast reveal context-dependent phenotypes and in trans effects on the protein regulatory network. *PLoS Genetics*, *17*(1), e1008761. https://doi.org/10.1371/journal.pgen.1008761

Baryshnikova, A., Costanzo, M., Dixon, S., Vizeacoumar, F. J., Myers, C. L., Andrews, B., & Boone, C. (2010a). Synthetic genetic array (SGA) analysis in Saccharomyces cerevisiae and Schizosaccharomyces pombe. *Methods in Enzymology*, *470*, 145–179. https://doi.org/10.1016/S0076-6879(10)70007-0

Baryshnikova, A., Costanzo, M., Kim, Y., Ding, H., Koh, J., Toufighi, K., Youn, J.-Y., Ou, J., San Luis, B.-J., Bandyopadhyay, S., Hibbs, M., Hess, D., Gingras, A.-C., Bader, G. D., Troyanskaya, O. G., Brown, G. W., Andrews, B., Boone, C., & Myers, C. L. (2010b). Quantitative analysis of fitness and genetic interactions in yeast on a genome scale. *Nature Methods*, *7*(12), 1017–1024. https://doi.org/10.1038/nmeth.1534

Chu, A. M., & Davis, R. W. (2008). High-Throughput Creation of a Whole-Genome Collection of Yeast Knockout Strains. In A. L. Osterman & S. Y. Gerdes (Eds), *Microbial Gene Essentiality: Protocols and Bioinformatics* (pp. 205–220). Humana Press. https://doi.org/10.1007/978-1-59745-321-9_14

Parker, S., Fraczek, M. G., Wu, J., Shamsah, S., Manousaki, A., Dungrattanalert, K., de Almeida, R. A., Estrada-Rivadeneyra, D., Omara, W., Delneri, D., & O'Keefe, R. T. (2017). A resource for functional profiling of noncoding RNA in the yeast Saccharomyces cerevisiae. *RNA (New York, N.Y.)*, *23*(8), 1166–1171. https://doi.org/10.1261/rna.061564.117
